# The paradox of neutral carbonate budgets on coral-dominated reefs

**DOI:** 10.64898/2026.05.11.724394

**Authors:** Elizabeth Cabrera-Rivera, Didier de Bakker, Ana L. Molina-Hernández, Francisco Medellín-Maldonado, Rodolfo Rioja-Nieto, Alexis E. Medina-Valmaseda, Esmeralda Pérez-Cervantes, Chris T. Perry, Lorenzo Álvarez-Filip

**Affiliations:** Biodiversity and Reef Conservation Laboratory, Unidad Académica de Sistemas Arrecifales, Instituto de Ciencias del Mar y Limnología, Universidad Nacional Autónoma de México, Puerto Morelos, México; Geography, Faculty of Environment, Science and Economy, University of Exeter, Exeter, UK; PIESACOM, UMDI-Sisal, Facultad de Ciencias, UNAM, Puerto de Abrigo s/n, C.P, 97356, Sisal, Yucatán, México; Laboratorio de Análisis Espacial de Zonas Costeras, Unidad Multidisciplinaria de Docencia e Investigación-Sisal, Facultad de Ciencias, Universidad Nacional Autónoma de México, Mérida, Yucatán, México

**Keywords:** stress-tolerant, sea urchins, parrotfish, reef accretion, bioerosion, thermal variability

## Abstract

Coral reefs deliver vital services via a complex three-dimensional framework sustained by the balance between calcium carbonate production and erosion, or the net carbonate budget state. In many tropical western Atlantic reefs, ecological decline has reduced carbonate production, yielding near-neutral or negative budgets. Yet some reefs retain high coral cover and, theoretically, should also have high net positive budgets, yet often show modest carbonate accumulation. We used the remote reef of Cayo Arenas in the Campeche Bank, Gulf of Mexico, to test whether in reefs under suboptimal (variable) environmental conditions, high coral production is offset by robust bioeroder communities, producing neutral budgets. At 14 sites, we quantified carbonate producers and bioeroders to estimate gross production, bioerosion, and net budget states. Despite relatively high live coral cover, mean net carbonate budgets were approximately neutral. Crucially, this neutrality arose not from depressed biological activity (as in degraded reefs) but from an active equilibrium: vigorous carbonate production coupled with substantial bioerosion. These reefs, therefore, represent a contemporary, functional reef state in net stasis. Distinguishing active-neutral from impoverishment-neutral regimes is critical for predicting reef trajectories under environmental change and for targeting management, although near-stasis emerging from high carbonate turnover can appear functionally intact yet operate with limited buffering capacity against net carbonate loss.

## Introduction

Biodiversity and ecosystem services associated with coral reefs largely depend on the reef structural framework, which is formed through the net biogenic accumulation of calcium carbonate (CaCO_3_) over ecological, geomorphic, and geological timescales from years to millennia (Perry et al., 2008). Reef formation and persistence is underpinned by two fundamental processes: the production and erosion of calcareous structures. The production of CaCO_3_ (gross production) provides the carbonate that builds and cements the reef framework, with scleractinian corals being the primary calcifiers (Cornwall et al., 2021). By contrast, erosional processes remove this material from the reef matrix. Parrotfish and sea urchins are often the primary source of biological erosion (or bioerosion) (Molina-Hernández & Álvarez-Filip, 2024), whilst physical disturbances can also strip large volumes of framework carbonate off the reef and/or drive reef-building. Resultant rates of gross production and bioerosion contribute to determining the amount of CaCO_3_ that remains in the reef structure, and the balance is often described as the calcium carbonate net budget or net production (Scoffin et al., 1980; Stearn et al., 1977). In this sense, the calcium carbonate net budget has become a tool for quantitatively estimating how gross production and bioerosion processes determine aspects such as habitat provision, coastal protection and conservation status (Lange et al., 2020).

There are three possible carbonate-dynamic scenarios on reefs (Kleypas et al., 2001; Perry, 2011; Perry et al., 2008). A positive net budget is characterised by high cover of reef-building corals whose CaCO_3_ accumulation supports framework construction and net vertical reef growth (scenario I, Fig. 1) and is generally considered optimal for tracking sea-level change and maintaining key reef functions (Perry et al., 2018, 2025). Conversely, a negative net budget arises when erosion exceeds gross production, producing net loss of CaCO_3_ (scenario III, Fig. 1); these states are typically associated with ecological degradation, low coral cover and dominance by slow⍰calcifying species (Estrada-Saldívar et al., 2021; Lange et al., 2022). Between these extremes is reef stasis, where gross production and erosion are balanced and framework accumulation is very slow or negligible. Importantly, this neutral condition can occur at distinct operating points: when both gross production and bioerosion are low (impoverished neutrality, scenario II, Fig. 1) or when both are high (active neutrality, scenario IV, Fig. 1). Impoverished neutrality reflects a functionally depleted, degraded system, whereas active neutrality denotes a balanced, structurally robust and functionally active reef. In comparison with impoverished neutrality, active neutrality is driven by high cover of key reef-building corals and a considerable abundance of high-rate bioeroding taxa such as parrotfish and sea urchins (Cabrera-Rivera et al., 2025; Randazzo-Eisemann et al., 2024).

**Figure 1.**
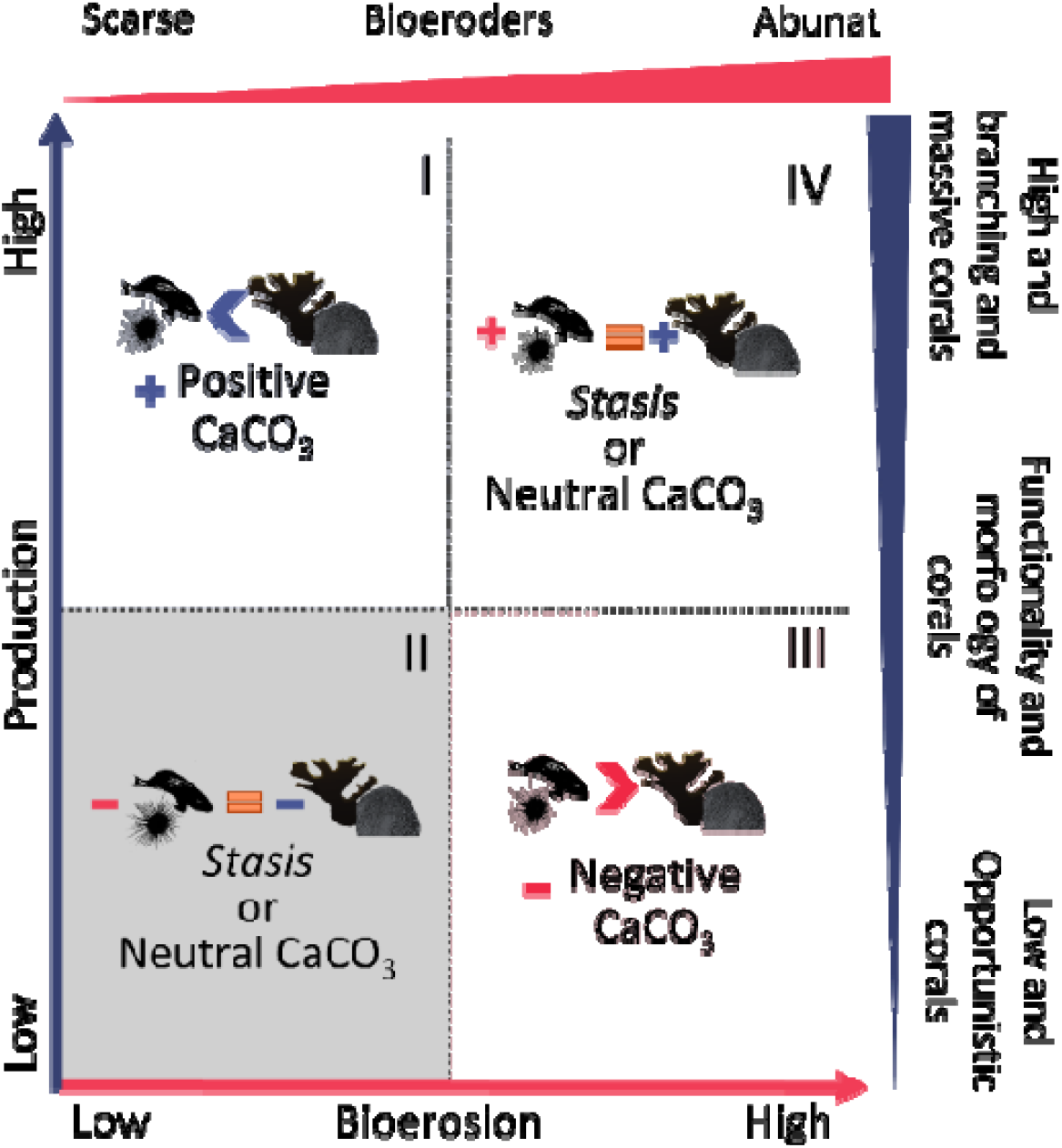
Conceptual model of four possible scenarios for net calcium carbonate net budgets, taking into account the ecological context. The right margin shows coral functionality and morphology, increasing from bottom to top. Encrusting secondary producers, such as crustose coralline algae (CCA), are included on the far bottom. The top margin indicates the presence or absence of bioeroders, with abundance increasing from left to right. Quadrant I exemplifies a positive, high-budget state, resulting from high gross production by high-functional corals, with bioerosion lower than gross production (bioerosion < gross production). Quadrant II shows a neutral net budget or impoverished neutrality; it is associated with a context of ecological degradation, where there is little contribution of CaCO_3_ from low-functional corals with CCA, and a low abundance of bioeroders (bioerosion = gross production). Quadrant III refers to a negative net budget caused by an increase in bioeroder abundance that outweighs the low gross production of corals with low reef functionality (bioerosion > gross production). Quadrant IV refers to an active neutrality state; associated with high ecological activity, where gross production by high-functionality corals is offset by intense bioerosion driven by a high abundance of bioeroders (bioerosion = gross production).

Many Tropical Western Atlantic reefs currently occupy impoverished⍰neutral states (scenario II, Fig. 1), following acute or chronic coral mortality from bleaching and disease, which have greatly reduced calcifier abundance (Alvarez-Filip et al., 2022; Courtney et al., 2022; Goreau and Hayes, 2024). Concurrent reductions in bioerosion — notably declines in sea urchin and parrotfish densities — have allowed reefs with low gross production to persist in stasis or slight accretion, a pattern common across the region (Molina-Hernández et al., 2020; Perry et al., 2018). By contrast, active neutrality is less examined despite being ecologically plausible, particularly in reefs developed under suboptimal conditions (e.g. turbid, eutrophic environments), which may characterise most of the world’s coral reefs (Morais et al., 2025). Assemblages in such settings are often dominated by massive and submassive, stress⍰tolerant corals that can sustain moderate–high carbonate production but rarely attain the extreme gross production of fast⍰growing branching taxa (Schoepf et al., 2023). Moreover, suboptimal conditions commonly reduce species⍰level skeletal extension and density relative to optimal settings, limiting community calcification (Carricart-Ganivet et al., 2012; Gutiérrez-Estrada et al., 2025; Morgan et al., 2016). Under these conditions, elevated yet structurally constrained carbonate production may be effectively offset by healthy and more active rates of bioerosion (Morris et al., 2022; Webb et al., 2017). Therefore, resulting in near-neutral carbonate budgets maintained through high process turnover rather than through functional collapse. This highlights the necessity of using local, contemporary measurements of gross production and bioerosion to characterise reef carbonate dynamics accurately.

To evaluate the operability of active neutrality (scenario IV, Fig. 1), we quantified gross production and bioerosion at Cayo Arenas. This small, relatively isolated reef system remains in good ecological condition, with high abundances of key reef-building corals and a healthy population of bioeroders (Fig. 2; Cabrera-Rivera et al., 2025; Frías-Vega et al., 2025). Nevertheless, Cayo Arenas lies in a region of highly variable environmental conditions, including elevated variability in sea surface temperature that reduces coral calcification (Carricart-Ganivet, 2004; Carricart-Gavinet & Merino, 2001; Sánchez-Pelcastre et al., 2023) We therefore hypothesise that Cayo Arenas exhibits active, rather than impoverished, neutrality driven by the co-occurrence of robust carbonate producers and active bioerosion under variable environmental conditions. To test this hypothesis, we conducted ecological, gross production and bioerosion process characterisations of sites across this system with the *ReefBudget* methodology (Perry & Lange, 2019), incorporating locally measured calcification rates for *Orbicella* spp. and extensive sea urchin surveys to account for this cryptic, nocturnal bioeroder (Cabrera-Rivera et al., 2025). We then place our results in the context of comprehensive Western Caribbean data to contextualise and discuss the implications of our findings.

**Figure 2.**
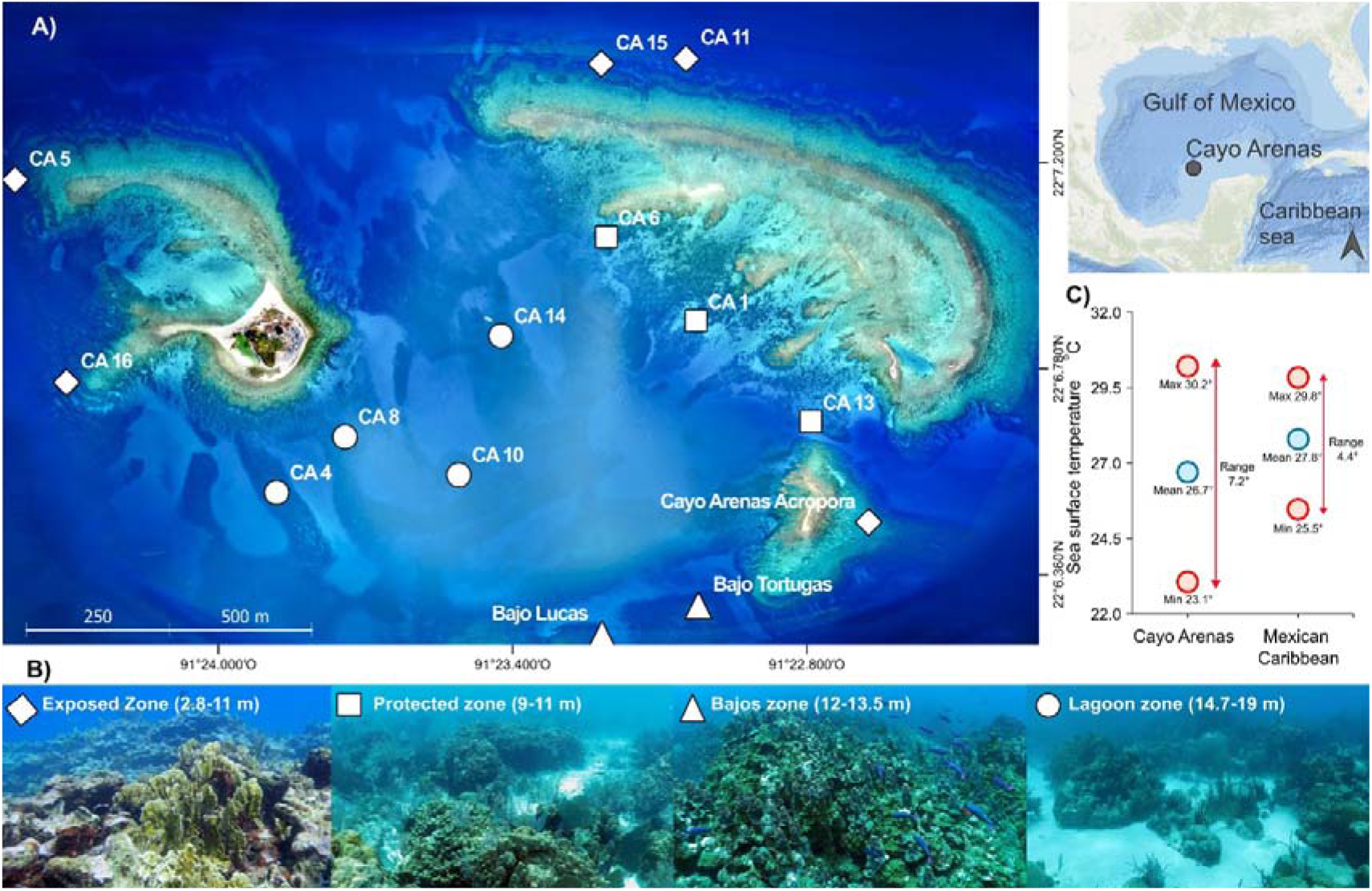
Location of Cayo Arenas and general information about the reef. Distribution of the fourteen sampled sites (A) and their classification into four reef zones (B). Also shown are the maximum, minimum, and average sea temperatures, as well as the range (°C) for Cayo Arenas and the Mexican Caribbean for comparation (C). Top right inset shows the positioning of Cayo Arenas in the Gulf of Mexico. The sea temperature data was obtained from Bio-ORACLE satellite data (Assis et al., 2024).

## Methods

### Study area

Cayo Arenas is a remote platform reef ∼200 km from the Yucatán coast, has an area of 10 km^2^ (Frías-Vega et al., 2025), and forms part of the Campeche Bank reef system in the southeast of the Gulf of Mexico (Fig. 2). It comprises a complex of reef structures that reach depths of up to 30 m and remain connected (Tunnel Jr. & Chávez, 2013). The underlying platform of the Campeche Bank is composed mainly of solidified carbonate materials (Logan et al., 1969) and exists in a region with highly variable environmental conditions (Molina-Hernandez et al *submitted*), such as high sea surface temperature (+7°C, Assis et al., 2024; Fig. 2). These result in low coral calcification and likely limit vertical reef growth (Blanchon & Perry, 2004; Carricart-Gavinet & Merino, 2001). The northeast structure of Cayo Arenas is the largest formation and has two emergent areas; the southeast structure is the smallest and has a small emerging area. These two structures receive the greatest impact from wind and wave energy (Fig. 2). The eastern structure, which has a lagoon on the windward side and a cay with vegetation, is in the most protected part of the system (Logan et al., 1969). Cayo Arenas reefs are dominated by a high abundance of large scleractinian massive corals, including *O. faveolata, O. annularis, Montastrea cavernosa, Colpophyllia natans*, and *Siderastrea* spp. (Jordán-Dahlgren, 2004; Perry et al., 2025).

### Data collection

No permissions were required for this research, as we did not collect or manipulate organisms, and the surveys were conducted before Cayo Arenas was decreed a Natural Protected Area in September 2025. For this study, 14 sites distributed across the four reef zones of Cayo Arenas (exposed, protected, bajos, and lagoon; Fig. 2) were surveyed in four expeditions between July 2017 and August 2024. Some sites (CA4, CA13, Bajo Tortugas, and CA1) were monitored for more than one year; however, because we did not examine temporal changes, we used only the most recent year with information for both producers and bioeroders. Due to the complexity of monitoring current intensity and reef depth, the number of transects per group of organisms varied across sites (see Tables S1 and S2; Cabrera-Rivera *submitted*).

At each site, a census was conducted using the *ReefBudget* V2 method for the Caribbean Sea (Perry & Lange, 2019) except at the deeper site CA10, where the point-intercept transect method was used (Table S1). For bioeroders, a belt transect method was used (10 x 1 m) for sea urchins and endolithic sponges, and 30 x 2 m (60 m^2^) for all non-cryptic reef fish species, with particular focus on parrotfishes given their functional relevance (Perry & Lange, 2019) . In the case of fish, the species, number of individuals, and fork length of each individual were recorded and classified by size (<5 cm, 8 cm, 15 cm and so on in 10 cm increments up to >120 cm). In addition to parrotfish, we also recorded the abundance and size of commercially important fish (groupers and snappers) and the main sea urchin predators (see Fig. S1 for details).

Furthermore, we conducted night-time monitoring to survey sea urchins at four sites (Bajo Tortugas, CA1, CA14, and CA15), as bioerosion rates by this group, based on daytime surveys, typically underestimate their abundance due to their diurnal cryptic lifestyle (Cabrera-Rivera et al., 2025; Courtney & Andersson, 2025). Each surveyed site represented one of the four zone reefs (Fig. 2). To make data collection with diving torches more efficient, the transect dimensions were modified to 20 x 0.5 m, while maintaining the sampling area (10 m^2^; Cabrera-Rivera et al., 2025). Surveys started at least 90 minutes after sunset. For logistical and safety reasons, it was not possible to survey all sites at night.

### Ecological characterization of Cayo Arenas

Although the primary focus of this study is on reef carbonate budgets, we conducted a broader ecological characterisation of Cayo Arenas derived from the same visual census data used throughout the study. We included metrics of live coral cover, the total absolute cover (% ± SE) of five coral groups (González-Barrios and Álvarez-Filip, 2018; Perry and Lange, 2019; Fig. S2, A), the density (ind m^-2^ ± SE) and/or biomass (kg 100 m^-2^ ± SE) of bioeroders, and the biomass of commercially important fishes as well as sea urchin predators (Fig. S3, A). These metrics were used solely to ecological context of carbonate budget, rather than as a standalone assessment of reef status. Fish biomass was estimated based on the size and abundance of species using the allometric equation and parameters from Froese and Pauly (2025). We estimated the population density of sea urchins (*Diadema, Echinometra* and *Eucidaris*) with data from night surveys.

We used the Mexican Caribbean as a reference to situate our findings in a broader ecological context. We use this region as a reference because it is well-studied, geographically closest, and most connected to the Campeche Bank (Jordán-Dahlgren, 2002; Jordan-Dahlgren & Rodriguez-Martinez, 2003). To interpret the impact of fishing on Cayo Arenas, the sites in the Mexican Caribbean were grouped according to the classification of Espinosa-Andrade et al (2020) as “Well-Managed Fisheries” and “Under-Managed Fisheries” (Table S4). Based on mean values obtained from the literature for the period 2017 - 2021 (Espinosa-Andrade et al., 2020; Molina-Hernández et al., 2020; Perry et al., 2018, 2025). Since sea urchins are not an important species in Caribbean fisheries, we do not distinguish between these areas and group all sites as the Mexican Caribbean.

### Reef Carbonate Budget Estimation

We estimated carbonate budgets by quantifying gross production and bioerosion using the *ReefBudget* v2 protocol (Perry and Lange, 2019). Net carbonate budget was expressed as the sum of all production minus total bioerosion, standardised to kg CaCO_3_ m^−2^ yr^−1^. Gross production was estimated primarily from field-derived benthic cover (including CCA) using point⍰intercept transects, combined with species-specific skeletal density and linear growth extension rates from the *ReefBudget* v2 database for the Gulf of Mexico and Caribbean (Perry and Lange, 2019). For the point⍰intercept method, a cm linear measurement was assigned to each recorded coral patch and other benthic element, after which the *ReefBudget* protocol was applied to compute site gross production.

To better reflect local conditions, we adjusted density and linear-growth values from the *ReefBudget* database via a geographic filter (Gulf of Mexico → Campeche Bank → Cayo Arenas; Table S3). This provided site and region-specific information for *Orbicella* spp. (*O. annularis* for Cayo Arenas; *O. faveolata* for the Gulf of Mexico). For other morphotypes (massive, submassive, digitate, branching), extension rates were taken from Gulf of Mexico data, and skeletal density from Caribbean regional averages (Table S3).

Because light attenuation affects calcification (Reynaud et al., 2007), we incorporated depth (light availability) effects on the dominant reef-builder *O. faveolata* using the Gutiérrez-Estrada et al (2025) model, which predicts calcification from the PAR attenuation coefficient (K_dPAR_). We used K_dPAR_ = 0.067 m^−1^ for Cayo Arenas (Assis et al., 2024) and mean sampling depths (Table S1) to estimate depth⍰adjusted calcification rates for *O. faveolata* at each site, constraining modeled calcification not to exceed observed averages for 5 – 10 m depths (0.85 g cm^−2^ yr^−1^; Sánchez-Pelcastre et al., 2023). These site⍰specific *O. faveolata* calcification coefficients were incorporated into the *ReefBudget* gross⍰production formulas. We applied depth adjustments only to *O. faveolata* because species⍰specific calcification– light relationships are not available for other taxa.

Bioerosion rates followed *ReefBudget* v2 parameters and equations (Perry and Lange, 2019). Parrotfish erosion was estimated from bite rate (bites day^−1^), fish size and life stage, bite volume (cm^3^), proportion of bites that remove substrate, and substrate density (g cm^−3^), yielding kg CaCO_3_ day^−1^ which was standardized to area and year. Endolithic sponge erosion used species⍰specific substrate area (m^2^) and published annual bioerosion rates (De Bakker et al., 2018), divided by transect area to give kg CaCO_3_ m^−2^ yr^−1^. Microbioerosion used the proportion of hard substrate available for microboring (excluding dead coral and algal mats) with published average annual rates (kg CaCO_3_ m^−2^ yr^−1^). Sea⍰urchin erosion was estimated at species level and test⍰size distributions on transects, using published regressions relating test size to erosion (g urchin^−1^ day^−1^; Perry and Lange, 2019), and converted to kg CaCO_3_ m^−2^ yr^−1^.

For sites where sea urchins were recorded only during daytime surveys, we applied day-night conversion factors derived from one site from each of the four reef zones where both diurnal and nocturnal surveys were conducted (see Table S1; Cabrera-Rivera et al., 2025). These zone⍰specific conversion factors were used except for Bajo Lucas, for which we applied nocturnal rates from the neighbouring Bajo Tortugas because conversion produced unrealistically high estimates. Total bioerosion per transect was the sum of all group contributions; only nocturnal sea⍰urchin estimates were used in the reported total bioerosion, while daytime rates were retained for other bioeroder groups.

We tested the influence of (1) using reduced, area⍰specific *O. faveolata* gross⍰production rates and (2) including diurnal versus nocturnal sea⍰urchin bioerosion across reef zones. Because the data violated the assumptions of normality and homoscedasticity, we used paired Wilcoxon signed⍰rank tests. For *O. faveolata*, we compared gross production (kg CaCO_3_ m^−2^ yr^−1^) from Gulf of Mexico literature values (Table S3) with calcification rates estimated from the Gutiérrez-Estrada et al (2025); and for sea⍰urchin erosion, daytime and nighttime rates were compared. All analyses were performed in R v. 2023.12.0 (R Core Team, 2023).

## Results

### Cayo Arenas is a reef in good ecological condition

Cayo Arenas has a high average (±SE) coral cover of 29.5 ± 1.4% (Fig. 3A). With the highest values at the bajos (34.8 ± 2.5%) and the protected zones (35.9 ± 3.4%). Whereas the lagoon (29.3 ± 2.4%) and the exposed zones (23.7 ± 2.2%) exhibit the lowest cover. Overall, the coral assemblage is dominated by massive taxa (19.6 ± 1.5%), most notably *Orbicella* spp. (14 ± 1.3%), particularly in zones protected from wave energy. In contrast, branching coral cover, associated with the highest calcification rates in the region, is low (0.78 ± 0.3%) in this system and is represented exclusively by *Acropora palmata*, which was restricted to two sites in the exposed zone. Non-reef-building coral species had the lowest cover (1 ± 0.3%). CCA cover was consistently high across reef zones (12.5 ± 1.4%), with peak values in the bajos (15.8 ± 2.3%) and exposed (20.4 ± 3.0%) zones.

**Figure 3.**
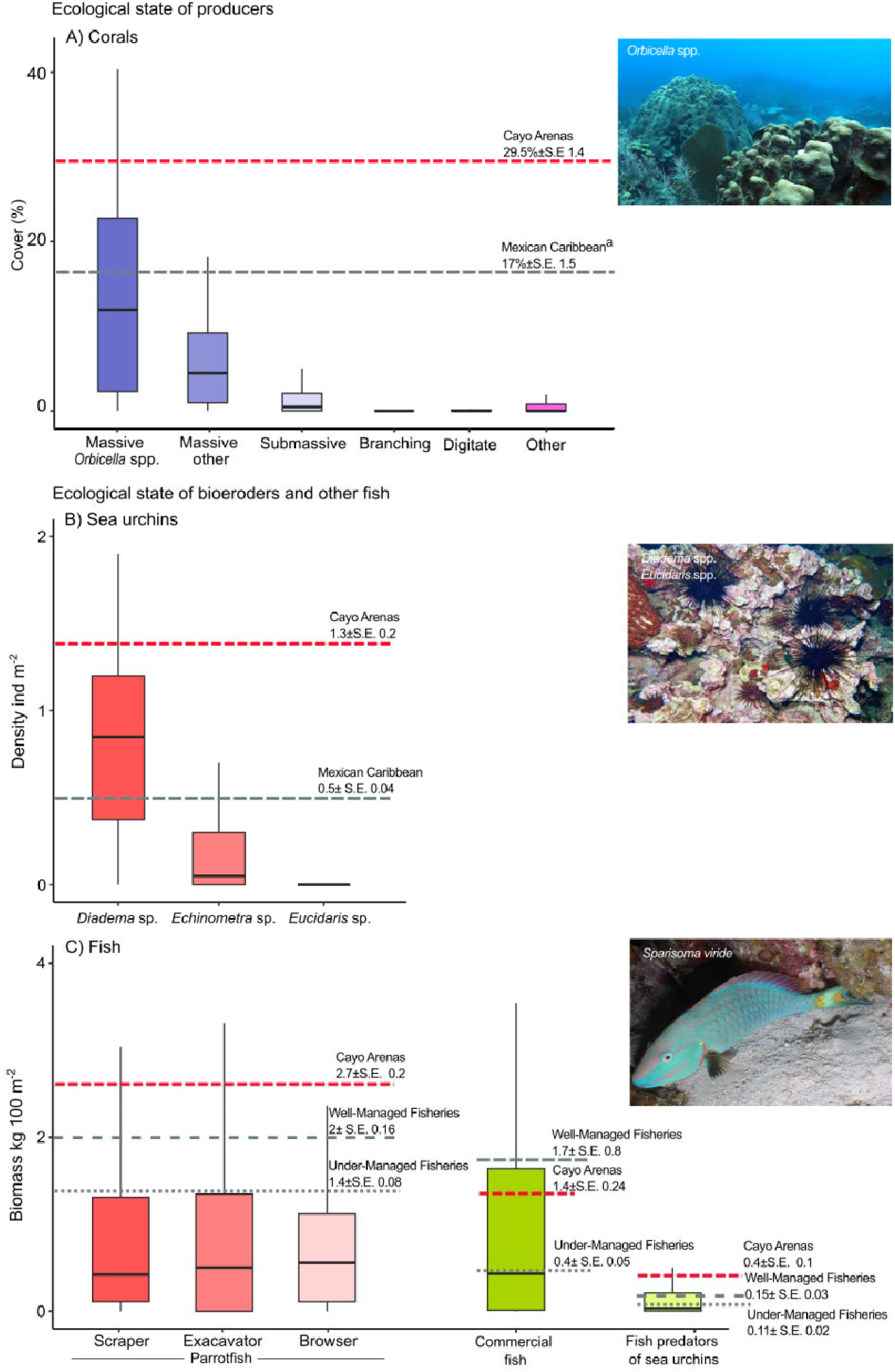
Ecological characterisation of Cayo Arenas reef. Comparisons were made with the Mexican Caribbean for coral cover and populations of eroding urchins and fish. The red dashed lines indicate the averages for Cayo Arenas. The grey dashed lines indicate the average for the Mexican Caribbean for coral cover and urchins, while for fish, they indicate the Well-Managed Fisheries in the Mexican Caribbean. The grey dotted lines indicate the Under-Management Fisheries in the Mexican Caribbean. The percentage of coral cover in the Mexican Caribbean was taken from Molina-Hernández et al (2020). Box plots indicate the interquartile range that contains 50% of the data. Lines within the boxes indicate the median. The lower and upper whiskers extend 1.5 of the interquartile range and show the variability of the data.

The average biomass of parrotfish in Cayo Arenas was 2.7 ± 0.2 kg 100 m^-2^ (Fig. 3C). The zone with the highest biomass was the exposed zone (4 ± 0.5 kg 100 m^-2^) and the zone with the lowest biomass was the bajos zone (1.3 ± 0.3 kg 100 m^-2^). The group of parrotfish with the highest biomass was the scraper, with 0.95 ± 0.1 kg 100 m^-2^, followed by the excavator, represented solely by the species *Sp. viride*, with 0.9 ± 0.1 kg 100 m^-2^. Then the browser, with 0.85 ± 0.1 kg 100 m^-2^. The average biomass of parrotfishes in Cayo Arenas was higher than that observed in the Caribbean, exceeding by ∼50% in the Under-Managed sites and by ∼25% in the Well-Managed sites (Fig. 3C).

The average biomass of commercially important fishes in Cayo Arenas was 1.4 ± 0.24 kg 100 m^-2^ (Fig. 3C). The protected zone had the highest biomass with 2.1 ± 0.9 kg 100 m^-2^, while the bajos zone had the lowest biomass with 1.1 ± 0.5 kg 100 m^-2^. The biomass of commercially important fishes observed at Cayo Arenas was slightly lower compared to that of Well-Managed Fisheries in the Caribbean (1.7 ± 0.8 kg 100 m^-2^), but considerably higher than that observed in Under-Managed Fisheries (0.4 ± 0.05 kg 100 m^-2^). Sea urchin predator fishes had higher valuers (0.4 ± 0.01 kg 100 m^-2^; Fig. 3C) than those observed in Well-Managed Fisheries (0.15 ± 0.03 kg 100 m^-2^) and Under-Managed Fisheries (0.11 ± 0.02 kg 100 m^-2^) in the Caribbean, suggesting a top-down control of sea urchin population. The exposed zone had the highest biomass of sea urchin predators (0.7 ± 0.3 kg 100 m^-2^) while the protected zone had the lowest biomass (0.2 ± 0.1 kg 100 m^-2^). In the case of eroding sea urchins, Cayo Arenas registered a population density of 1.3 ± 0.2 ind m^-2^ (Fig. 3B). The zone with the highest sea urchin density was the bajos (2.8 ± 0.95 ind m^-2^), while the zone with the lowest density was the lagoon (0.02 ± 0.02 ind m^-2^). *D. antillarum* had the highest population (1 ± 0.2 ind m^-2^), followed by the *Echinometra viridis* (0.2 ± 0.6 ind m^-2^), *E. lucunter* (0.01 ± 0.2 ind m^-2^) and finally *Eucidaris tribuloides* (0.02 ± 0.05 ind m^-2^). *D. antillarum* was also highly associated with the bajos zone, having the highest record of this species (2.5 ± 0.9 ind m^-2^). Sea urchin population in the Mexican Caribbean is less than half that of Cayo Arenas (Fig. 3B).

### Production and bioerosion rates are in a relative equilibrium

Cayo Arenas gross production rate averages 3.5 ± 0.3 kg CaCO_3_ m^-2^ yr^-1^, which is attributed almost entirely to the contribution of corals, with CCA accounting for only ∼1% (0.05 ± 0.0 kg CaCO_3_ m^-2^ yr^-1^). The species with the highest contribution to gross production is *O. faveolata* (1 ± 0.1 kg CaCO_3_ m^-2^ yr^-1^, Fig. S2B). At the same time, the mean bioerosion rate for the system was only slightly lower than the gross production (2.4 ± 0.2 kg CaCO_3_ m^-2^ yr^-1^). Parrotfish made the greatest contribution to net bioerosion (1.5 ± 0.2 kg CaCO_3_ m^-2^ yr^-1^), primarily driven by *Sp. viride* and *Sc. vetula* in the exposed and protected zones (Table S2 and S5). Yet sea urchins also exhibited a high bioerosion rate (1 ± 0.2 kg CaCO_3_ m^-2^ yr^-1^), particularly by *D. antillarum* in reefs with higher coral complexity in the bajos zone (Fig. S5, Table S6 and S7). The contributions of endolithic sponges (0.08 ± 0.08 kg CaCO_3_ m^-2^ yr^-1^) and microbioerosion (0.1 ± 0.0 kg CaCO_3_ m^-2^ yr^-1^) were modest to the overall bioerosion (Fig. S3C, Table S2, S9 and S10).

Overall, the net budget was close to stasis (0.8 ± 0.3 kg CaCO_3_ m^-2^ yr^-1^), as high bioerosion rates closely counterbalanced gross CaCO_3_ production. The zones with the highest net budgets were the protected and lagoon zones (3.1 ± 0.6 and 1.3 ± 0.4 kg CaCO_3_ m^-2^ yr^-1^, respectively), where the processes of bioerosion seemed to be less active (Fig. S4 and S5, Table S2). Whereas the bajos and exposed zones had the lowest net budgets (0.9 ± 0.7 and -1.1 ± 0.3 kg CaCO_3_ m^-2^ yr^-1^, respectively), largely owing to the high level of parrotfish and sea urchin bioerosion observed on those sites (Fig. S4 and S5, Table S2). Although there is a high level of variability at the transect level (dots in Fig. 4 and S4), 90% of sites have a net budget state relatively low (from -2 to 2 kg CaCO_3_ m^-2^ yr^-1^; Table S2), also it is worth mentioning that the transects with the highest level of net erosion were in the exposed zone, while highly net positive transects occurred mainly in the protected zone.

**Figure 4.**
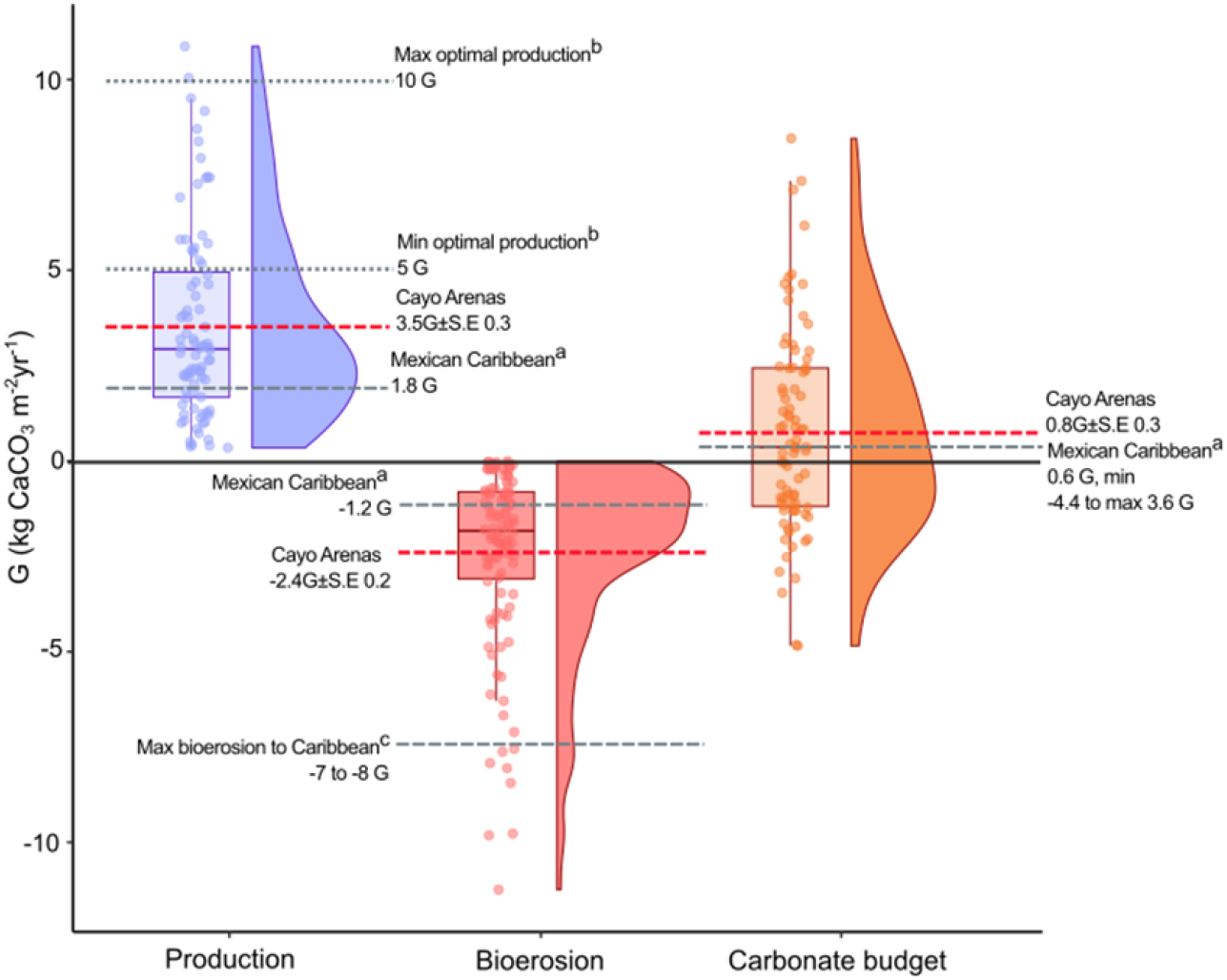
Gross production rates, bioerosion, and carbonate net budget of Cayo Arenas. Averages for Cayo Arenas are shown (dashed red lines). Points are transect-level values. References were included to compare the Mexican Caribbean (Molina-Hernández et al. (2020)^a^; dashed grey lines), the historical maximum bioerosion estimate for all Caribbean (Perry et al. (2014)^c^; dashed grey lines), and optimal gross production (5 to 10 kg CaCO_3_ m^-2^ yr^-1^ = G) estimated and considered for tropical forereefs (Vecsei (2004)^b^; dotted grey lines). Box plots indicate the interquartile range containing 50% of the data. Lines within the boxes indicate the median. The lower and upper whiskers extend 1.5 of the interquartile range and show the variability in the data. Distribution diagrams are shown above the box plots; each point represents the value for a given transect. Violin plots show probability density.

## Discussion

Our findings confirm that scenarios with neutral net budgets, characterised by net carbonate production values close to zero, do not necessarily indicate degraded habitats with low coral cover and depleted fish stocks (scenario IV, Fig. 1). Instead, we found that Cayo Arenas functions as a robust reef system (*sensu* Brandl et al., 2019), displaying high cover of key reef-building corals and the presence of herbivores (e.g., parrotfish and *D. antillarum*) along with high biomass levels of high-trophic-level species (e.g., groupers, snappers) comparable to those in Well-Managed Fisheriess (Fig. 3C). Here, production stasis appears driven not by ecological collapse but by relatively low calcification rates of dominant coral species—likely a consequence of suboptimal environmental conditions, rather than by reduced coral abundance (Fig. 1; Cruz-Piñón et al., 2003; Sánchez-Pelcastre et al., 2023). Concurrently, high abundances of parrotfish and sea urchins indicate strong potential for removal of framework carbonate and sediment; the elevated sea urchin signal is particularly apparent at night, when individuals shelter by day in complex reef structure (Cabrera-Rivera et al., 2025; Fig. S5, Table S2 and S6). Thus, in functionally robust reefs such as Cayo Arenas, ecological interactions can sustain a modest net-neutral budget driven by the coexistence of vigorous carbonate production and active bioerosion, rather than by loss of ecological functionality.

Cayo Arenas can be classified as an extreme reef (*sensu* Schoepf et al., 2023) because it experiences large intra⍰annual variability in sea temperature, nutrient concentrations and other environmental parameters relative to historically defined “optimal” ranges for coral growth (Fig. 2; Molina-Hernández et al., *submitted*). Such variability exposes corals to recurrent stress yet fosters assemblages dominated by stress⍰tolerant taxa capable of maintaining structural integrity, relatively high coral cover (>30%) and geo⍰ecological functions. These massive species typically have lower calcification rates than fast⍰growing acroporids found in more ‘optimal’ settings (Buckingham et al., 2022), but their morphology and biological traits (thick tissue and reduced skeletal density) confer resilience to environmental fluctuations (van Woesik et al., 2012). The *Orbicella* complex, a historical reef builder throughout the Holocene in the western Atlantic (Toth et al., 2019), contributes substantially to modern reef CaCO_3_ production (González-Barrios and Álvarez-Filip, 2018; Gutiérrez-Estrada et al., 2025), highlighting its ecological importance and adaptive capacity. High *Orbicella* cover may reflect allocation of calcification to extended skeleton growth despite porous skeletons (Carricart-Ganivet, 2004; Carricart-Gavinet and Merino, 2001; Cruz-Piñón et al., 2003). Furthermore, colony⍰level plasticity in physiology and morphology enables these species to persist under variable light, depth, temperature and turbidity (Carricart-Ganivet, 2004; Freitas et al., 2019)), with direct consequences for community composition and contribution to reef carbonate construction.

Gross carbonate production at Cayo Arenas was below the reported optimal regional range of < 5-10 kg CaCO_3_ m^-2^ yr^-1^ (Vecsei, 2004), despite exhibiting high cover of reef-building coral species. This reflects the dominance of *Orbicella faveolata*, a massive reef-building coral whose calcification rates are lower than those of fast-growing branching taxa, particularly under seasonally temperature-stressed conditions. The low gross production values were related to the calcification model of *O. faveolata* to estimate its production (see methods and Fig. S5), and being the dominant species, it consequently reduced the gross production of the study area. Low gross production rates under suboptimal conditions are consistent with extreme reefs (Schoepf et al., 2023) experiencing extreme high and low temperatures or high turbidity levels (Céspedes-Rodríguez and Londoño-Cruz, 2021; Manzello et al., 2018; Randi et al., 2021). At greater depths and under turbid conditions, calcification decreases as light availability does (Baker & Weber, 1975; Gutiérrez-Estrada et al., 2025; Morgan et al., 2020). Long-term exposure to temperature variability has likely made corals at Cayo Arenas more resilient than those in the Mexican Caribbean. Currently, this is important because coral species that make up the majority of Cayo Arenas are highly susceptible to diseases that have caused significant mortality in the Caribbean Sea (Alvarez-Filip et al., 2022; Williams et al., 2020).

A notable result of this study is that sea urchins contributed nearly half of the total bioerosion at Cayo Arenas, a contrast to many Caribbean reefs still recovering from the 1980s *Diadema* die⍰off (Lessios, 2016), and consistent with reports of robust sea urchin populations in other reefs of the Gulf of Mexico (e.g. Veracruz, Banco Stetson, Flower Gardens; Johnston et al., 2021; Morales-Quijano et al., 2017; Nuttal et al., 2020). The sea urchin bioerosion rates throughout Cayo Arenas were significantly higher during nighttime surveys although their populations were also quite abundant during the daytime (Fig. S5B; Cabrera-Rivera et al., 2025). While high local bioerosion might signal an imbalance driven by excessive sea urchin density, often linked to overfishing and loss of predators (Carreiro-Silva and McClanahan, 2001), this appears unlikely here. The fact that fish biomass at Cayo Arenas is comparable to that of Well Managed Fisheries in the Caribbean (Fig. 3C) and the availability of structurally complex refuges (Cabrera-Rivera et al., 2025) suggest a currently stable and controlled urchin population. Nonetheless, given the high erosion rates in some areas of Cayo Arenas and elsewhere in the Gulf of Mexico, intensified fishing, which removes top-down control, could rapidly shift the system toward highly erosive states observed elsewhere.

Parrotfishes were the primary bioeroders at Cayo Arenas, consistent with their well-documented role as the dominant bioeroding group in Atlantic and other reef systems (e.g., Glynn and Manzello, 2015). Among the most abundant species were species *Sp. viride* and *Sc. vetula*, which, in addition to being the two species with the highest eroding potential in the Atlantic Ocean, are also the main sediment producers (Molina-Hernández and Álvarez-Filip, 2024). This is important because these species are currently highly vulnerable to fishing, which could compromise the reef’s sediment supply (Molina-Hernández and Álvarez-Filip, 2024). Parrotfish bioerosion at Cayo Arenas fell within values reported for the Campeche Bank (∼0.5–4 kg CaCO3 m^−2^ yr^−1^; Randazzo-Eisemann et al., 2024) but exceeded rates reported for nearshore reefs (Mexican Caribbean, Veracruz; 0.4 – 0.7 kg CaCO_3_ m^−2^ yr^−1^; Molina-Hernández et al., 2020; Perry et al., 2025). By contrast, microbioerosion and endolithic sponge erosion made only minor contributions to the budget, although their rates were within ranges reported elsewhere (e.g. Perry et al., 2025). This differs from some studies in which endolithic sponges play larger roles, often when sea urchins are absent, underscoring context⍰dependent shifts among bioeroder groups (de Bakker et al., 2019; Kuffner et al., 2019a).

Overall, Cayo Arenas exhibits near⍰neutral net carbonate budgets despite high abundances of carbonate producers due to local environmental constraints that limit gross production, and healthy populations of parrotfishes and sea urchins drive relatively high bioerosion (Fig. 3, 4 and S5, Table S2). This exemplifies active neutrality (scenario IV, Fig. 1), contrasting with impoverished neutrality at many Caribbean sites resulting from the deterioration of the coral community (Cornwall et al., 2021; Perry et al., 2025), and decreased bioerosion rates (e.g., Molina-Hernández et al., 2020; Perry et al., 2014). Remote reefs with strongly positive budgets typically report much higher production rates that outpace erosion (Cornwall et al., 2021; Lange et al., 2022; Morgan and Kench, 2016; Perry et al., 2015), but those estimates often rely on regional averages and do not account for depth⍰related or site⍰specific reductions in calcification. In our study, adjusting *O. faveolata* calcification to local conditions and including nocturnal urchin bioerosion lowered net estimates. This illustrated how local, contemporary measurements of calcification and bioerosion are essential for accurate carbonate⍰budget assessments and for predicting reef responses to increasing environmental threats. We therefore emphasise the importance of continuing to generate local information on calcification and bioerosion and including it in the estimation of carbonate net budgets, as this will allow us to gain more accurate insights into reefs’ adaptive responses to growing threats, such as global warming.

There are two relevant implications to consider for sites, such as Cayo Arenas, that have relatively good ecological conditions but near-neutral carbonate net budgets. First, it is likely that carbonate net budgets at this site is highly sensitive to drastic changes in some components of the system. For example, if a lethal disease outbreak or a mass bleaching causes widespread coral mortality in susceptible species (e.g., Alvarez-Filip et al., 2022), erosion rates could increase disproportionately. This would result not only from the loss of living tissue and the associated decline in gross carbonate production, but also from the exposure of skeletons of dominant massive corals such as *Orbicella faveolata*, whose skeletal traits render them highly susceptible to intense bioerosion (Kuffner et al., 2019b; Molina‐Hernández et al., 2022). In the presence of robust communities of external bioeroders, such a shift could rapidly drive the system into a strongly net-negative carbonate state, with implications for rapid framework degradation. Secondly, in the context of rapid anthropogenic climate change, the rates of vertical accretion at Cayo Arenas are nowhere near projected rates of sea-level rise (Perry et al., 2025). Consequently, even apparently resilient reefs will lag in long⍰term growth and likely will fail to keep pace with sea-level rise (Cornwall et al., 2023; Perry et al., 2025). This may lead to alterations in reef ecology, particularly in shallow and deep reef assemblages, which are now characterised by reduced gross carbonate production. Preserving the ecological integrity of these remote reefs should therefore be a priority in the coming years, as reefs such as Cayo Arenas remain largely unexplored biological reservoirs. Furthermore, they could function as potential refuges for coral restoration and reef biodiversity.

## Data accessibility

All data are provided as supplementary data (Cabrera-Rivera *et al., submitted*)

## Acknowledgments

We thank the Caribbean Kraken team and Manuel Victoria for all the support in sampling during the expedition to Cayo Arenas. This research was funded by the Programa de Apoyo a Proyectos de Investigación e Innovación Tecnológica, UNAM number IG201323. E. C-R received a master’s scholarship from the Secretaría de Ciencia, Humanidades, Tecnología e Innovación (SECIHTI, 1249631).

