## supporting material for "The paradox of neutral carbonate budgets on coral-dominated reefs"

**Supplementary material**

**Figure S1**

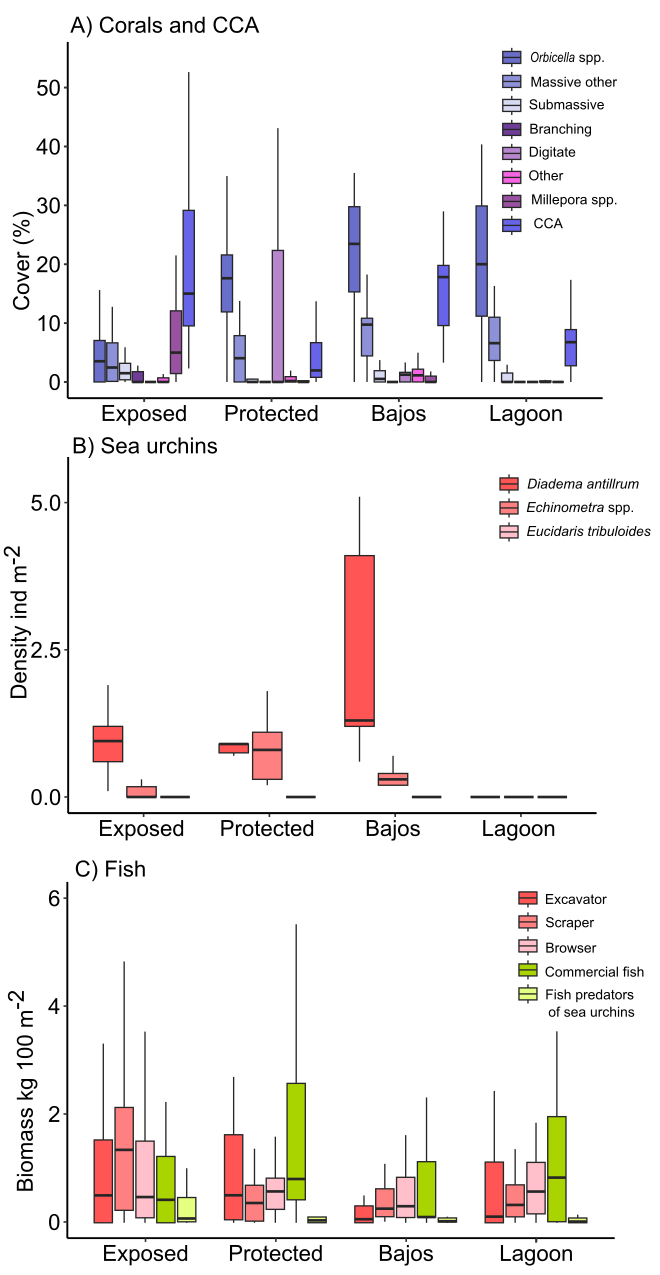

**Figure S1**. Ecological characterization at four reef zones of Cayo Arenas. For commercially important fish (groupers and snappers), we considered: *Cephalopholis cruentata, C. fulva, Epinephelus adscensionis, E. guttatus, E. striatus, Mycteroperca bonaci, M. interstitialis, M. tigris, M. venenosa, Lutjanus analis, L. apodus, L. cyanopterus, L. griseus, L. jocu, L. mahogoni, L. synagris and Ocyurus chrysurus (sensu* Lang et al., 2010)*.* Sea urchin predators included: *Balistes vetula, Bodianus rufus, Calamus calamus, Canthidermis sufflamen, Canthigaster rostrata, Diodon hystrix, Halichoeres bivittatus, H. garnoti, H. radiatus, Lactophrys bicaudalis* and *Thalassoma bifasciatum* (sensu Behrents et al., 1984, Randall et al., 1964, Rodríguez-Barreras et al., 2015.

**Figure S2**

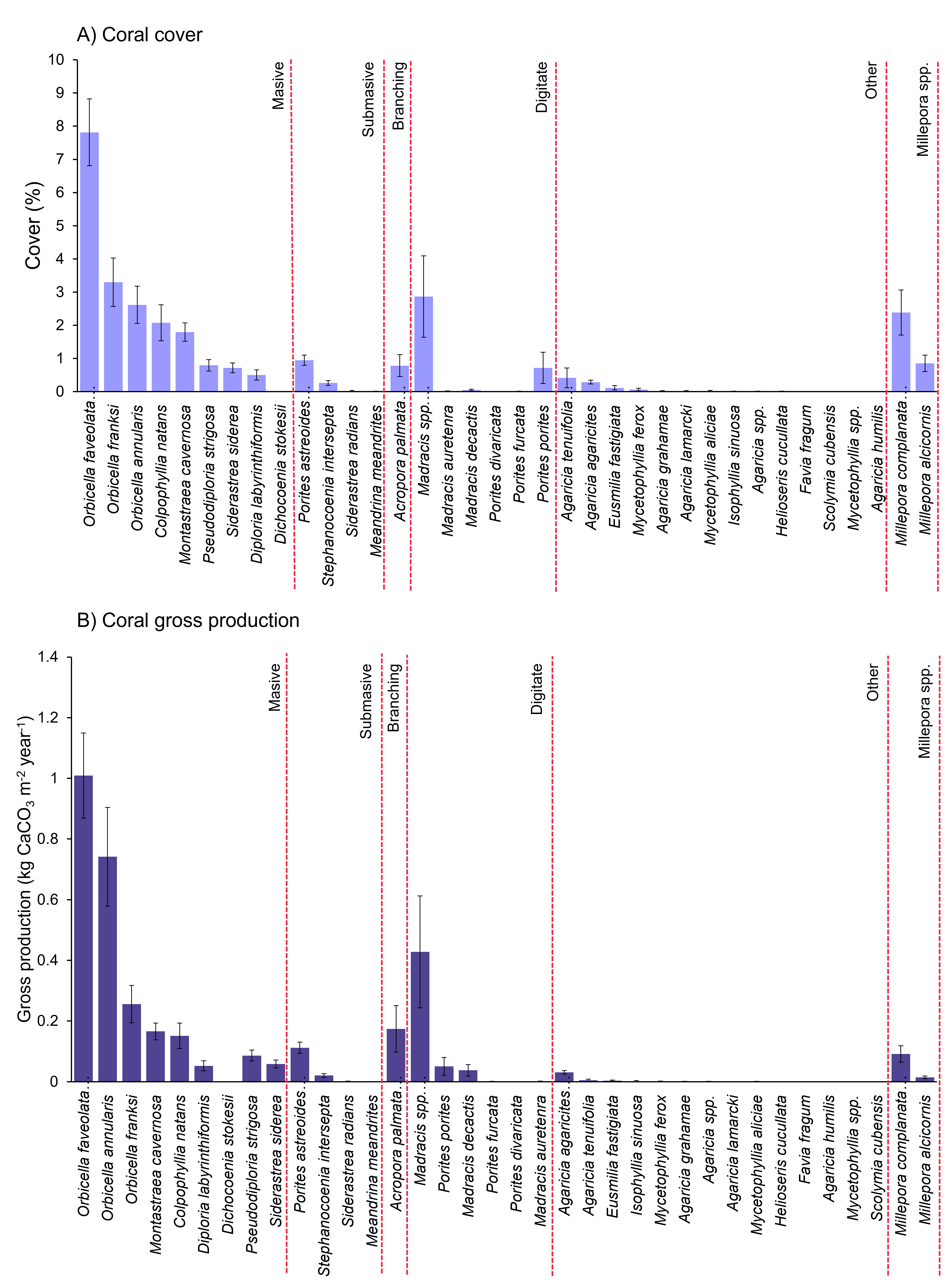

**Figure S2.** Cover and gross production of coral species in Cayo Arenas.

**Figure S3**

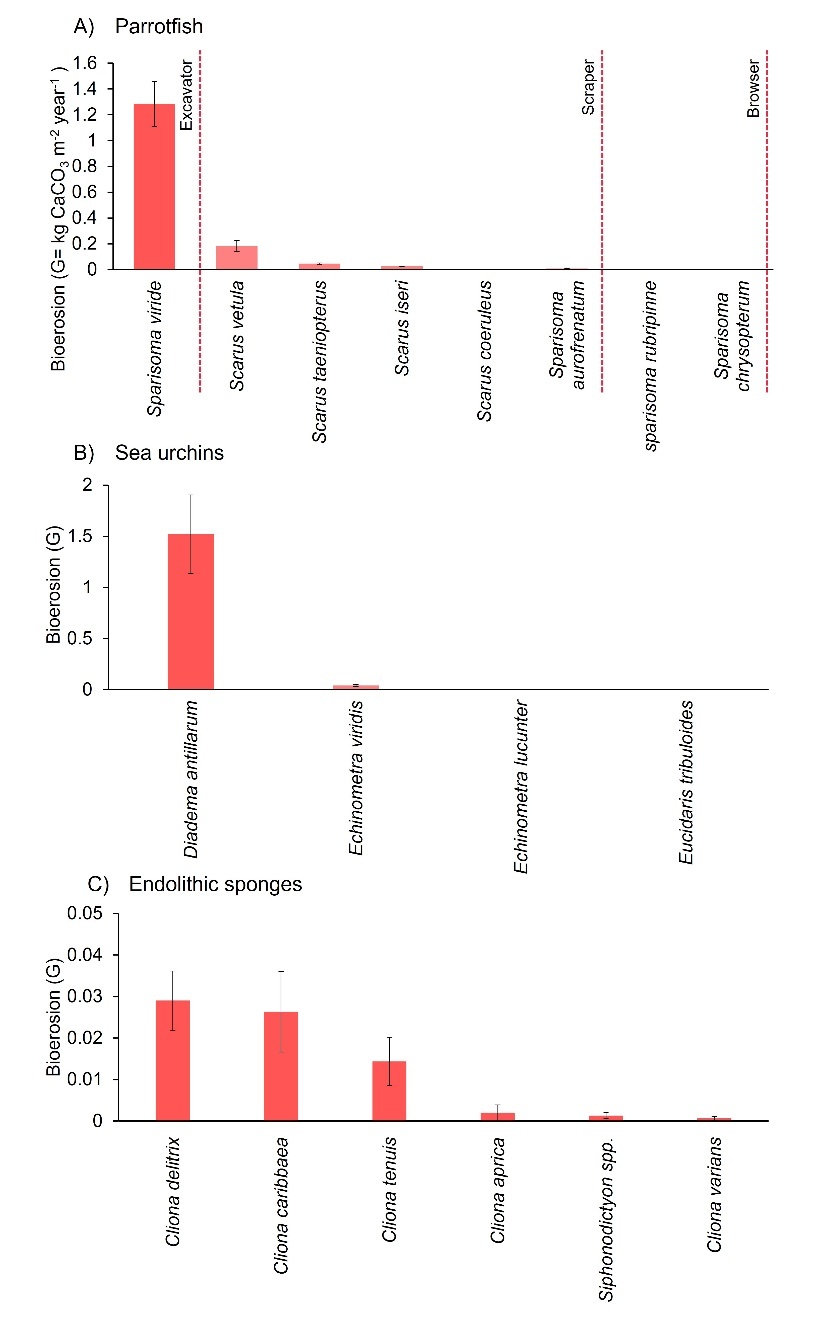

**Figure S3.** Bioeroders at Cayo Arenas and their bioerosion rates.

**Figure S4**

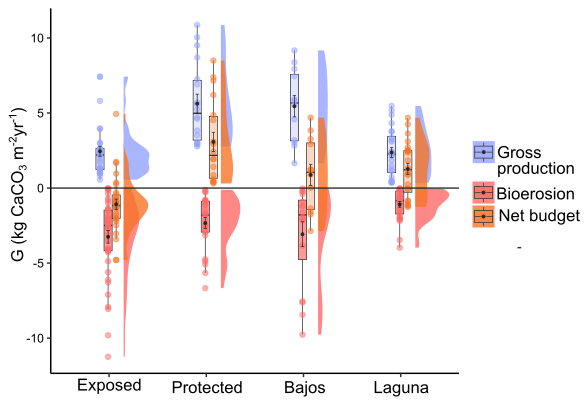

**Figure S4**. Gross production, bioerosion and net budget rates in four reef zones of Cayo Arenas.

**Figure S5.**

*Adjusted rates by using more reliable data on Orbicella faveolata and sea urchins.*

We found significant differences between the two methods used to estimate *O. faveolata* CaCO_3_ (Fig. S5). This is when rates were estimated using the Gulf of Mexico parameters versus using the calcification model that considers local rates and accounts for the depth effect (see methods). Gross production rates obtained using the calcification model are lower across all zones. The amount of *O. faveolata* gross production rates fell from 1.7 ± 0.2 to 1 ± 0.1 kg CaCO_3_ m^-2^ yr^-1^ (V= 0, p<0.05). However, the proportion of *O. faveolata* production in each reef zone was the same in both methods (Fig. S5, A).

Similarly, sea urchin bioerosion rates throughout Cayo Arenas were deemed to be significantly higher during nighttime surveys (Fig. S5, B). Sea urchin bioerosion goes from 0.4 ± 0.1 kg CaCO_3_ m^-2^ yr^-1^ at daylight to 1 ± 0.2 kg CaCO_3_ m^-2^ yr^-1^ (V= 1756.5, p<0.05) at night. We observed a trend showing that the bajos and protected zones had the highest O. faveolata gross production and nocturnal sea urchin bioerosion rates, while the other two zones had the lowest rates for both.

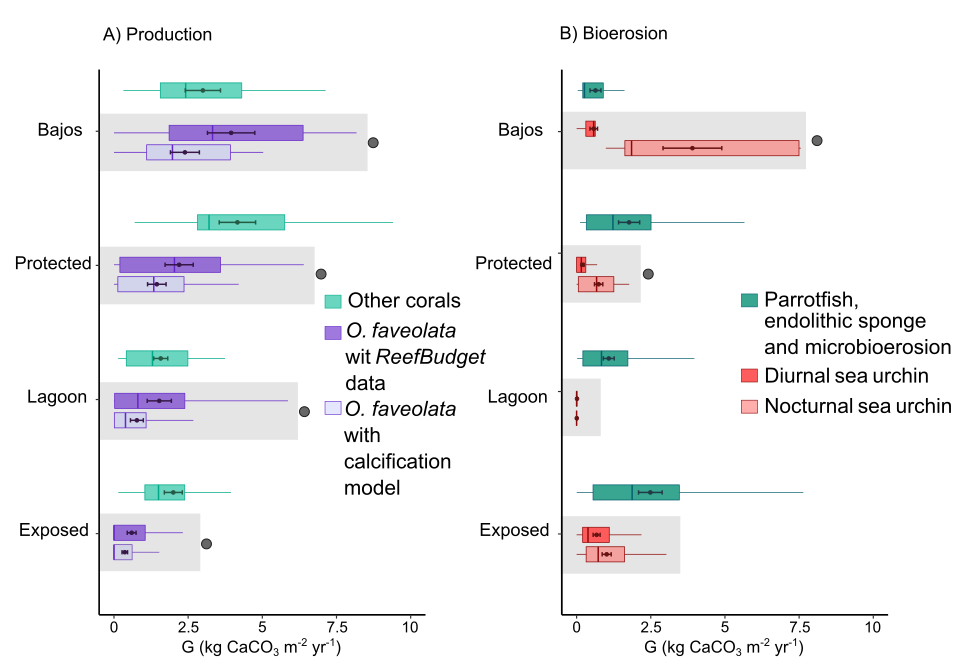

**Figure S5.** Comparison of *Orbicella faveolata* production rates and sea urchin bioerosion among Cayo Arenas reef zones. Overall and adjusted data. A) Calcium carbonate production for each reef zone is shown. Green indicates production by all corals (see Fig. S2B) except *O. faveolata*. The shaded area compares *O. faveolata* production estimated from depth and K_dPAR_ data with the rate estimated from the Gulf of Mexico data. Significant differences were observed between the two production types within each zone (circle, p<0.05). B) Bioerosion for each reef zone is shown, where green indicates bioerosion by all groups (parrotfish, endolithic sponges, and microbioerosion) except urchins. The shaded area compares diurnal and nocturnal urchin bioerosion. Black dots at the top of the shaded area indicate significant changes between samples within each zone (p<0.05). Box plots indicate the interquartile range containing 50% of the data. Lines within the boxes indicate the median. The lower and upper whiskers extend 1.5 times the interquartile range and show the variability in the data.

**Table S1.** Information about 14 sites monitored in Cayo Arenas and values used to calculate *Orbicella faveolata* gross production and nighttime sea urchin bioerosion.

| **Site** | **Reef zone** | **Year** | **Mean survey depth (m)** | **K_dPAR_ Cayo Arenas** | **Irradiance benthos** | ***Orbicella faveolata* calcification***  **(g cm^2^ yr^-1^)** | **Conversion factor to estimate nightime sea urcihin bioerosion** |
| --- | --- | --- | --- | --- | --- | --- | --- |
| Cayo Arenas 11 (CA11) | Exposed | 2017 | 10 | 0.077 | 46.36 | 0.855 | 1.47 |
| Cayo Arenas 15 (CA15) |  | 2023 | 8.8 |  | 50.84 | 0.82 |  |
| Cayo Acropora |  | 2021 | 2.8 |  | 80.94 | 0.38 |  |
| Cayo Arenas 5 (CA5) |  | 2017 | 8 |  | 54.06 | 0.79 |  |
| Cayo Arenas 16 (CA16) |  | 2023 | 12.4 |  | 38.55 | 0.83 |  |
| Cayo Arenas 6 (CA6) | Protected | 2017 | 9.9 |  | 46.72 | 0.854** | 3.59 |
| Cayo Arenas 1 (CA1) |  | 2023 | 9.9 |  | 46.84 | 0.853** |  |
| Cayo Arenas 13 (CA13) |  | 2017 | 11 |  | 42.93 | 0.856** |  |
| Bajo Tortugas | Bajos | 2023 | 14.4 |  | 32.97 | 0.76 | 13.36*** |
| Bajo Lucas |  | 2021 | 13.4 |  | 35.63 | 0.80 |  |
| Cayo Arenas 14 (CA14) | Lagoon | 2023 | 19 |  | 23.21 | 0.59 | 0 |
| Cayo Arenas 8 (CA8) |  | 2017 | 14.7 |  | 32.30 | 0.75 |  |
| Cayo Arenas 4 (CA4) |  | 2023 | 19.4 |  | 22.51 | 0.57 |  |
| Cayo Arenas 10 (CA10) |  | 2021 | 19.3 |  | 22.77 | 0.58 |  |

*Local calcification values used to substitute at formula gross production of *O. faveolata* at ReefBudget method (Perry and Lange, 2019). **Published mean calcification rate of *O. faveolata* for Cayo Arenas (0.85 g cm^2^ yr^-1^, Sánchez-Pelcastre et al., 2023). ***The bajos conversion factor was not applied to estimate nighttime bioerosion Bajo Lucas, as it increased the estimated nighttime bioerosion rate at this zone.

**Table S2**. Mean rates of gross production, bioerosión and carbonate net budget in Cayo Arenas.

| **Site** | **Reef zone** | **#**  **Benthos transec-**  **ts** | **X̅**  **Gross Produc-**  **tión (G)** | **X̅ Microbio-erosion (G)** | **#**  **Endolithic sponges transec-**  **ts** | **X̅ Endolithic sponges bioero-**  **sión (G)** | **#**  **Parrotfish transec**  **ts** | **X̅**  **Parrotfish bioero-**  **sion (G)** | **#**  **Sea urchins transec-**  **ts** | **X̅**  **Sea urchins bioero-**  **sion (G)** | **#**  **Nighttime sea urchins transec-**  **ts** | **X̅**  **Nighttime sea urchins bioerosion (G)*** | **∑ Total bioero-**  **sión (G)** | **Net budget CaCO_3_ (G)** |
| --- | --- | --- | --- | --- | --- | --- | --- | --- | --- | --- | --- | --- | --- | --- |
| CA11 | Exposed | 6 | 2.210 | 0.118 | 6 | 0.04 | 8 | 1.229 | 6 | 1.376 | NA | 2.026 | 3.413 | -1.203 |
| CA15* |  | 6 | 3.345 | 0.118 | 6 | 0.071 | 8 | 3.899 | 6 | 1.086 | 7 | 1.600 | 5.688 | -2.343 |
| Cayo Acropora |  | 6 | 3.110 | 0.069 | 6 | 0.169 | 8 | 1.874 | 6 | 0.214 | NA | 0.316 | 2.428 | 0.682 |
| CA5 |  | 6 | 1.746 | 0.155 | 6 | 0.001 | 9 | 2.264 | 6 | 0.453 | NA | 0.667 | 3.088 | -1.342 |
| CA16 |  | 6 | 1.819 | 0.116 | 6 | 0.018 | 8 | 2.531 | 6 | 0.245 | NA | 0.360 | 3.025 | -1.205 |
| CA6 | Protected | 6 | 4.514 | 0.160 | 6 | 0.033 | 10 | 1.675 | 6 | 0.176 | NA | 0.630 | 2.498 | 2.015 |
| CA1 |  | 6 | 5.641 | 0.182 | 6 | 0.016 | 8 | 1.344 | 6 | 0.331 | 7 | 1.188 | 2.729 | 2.912 |
| CA13 |  | 6 | 6.714 | 0.127 | 6 | 0.000 | 7 | 1.948 | 6 | 0.094 | NA | 0.337 | 2.412 | 4.302 |
| Bajo  Tortugas | Bajos | 6 | 4.680 | 0.092 | 6 | 0.059 | 8 | 0.490 | 6 | 0.292 | 5 | 3.903 | 4.545 | 0.135 |
| Bajo  Lucas |  | 6 | 6.231 | 0.113 | 6 | 0.052 | 8 | 0.546 | 6 | 0.864 | NA | 3.903 | 4.613 | 1.617 |
| CA14 | Lagoon | 6 | 3.175 | 0.107 | 6 | 0.024 | 8 | 0.433 | 6 | 0.000 | 6 | 0.021 | 0.585 | 2.590 |
| CA8 |  | 6 | 3.435 | 0.091 | 6 | 0.005 | 7 | 1.137 | 6 | 0.021 | NA | 0.000 | 1.233 | 2.202 |
| CA4 |  | 6 | 2.423 | 0.035 | 6 | 0.03 | 7 | 1.192 | 6 | 0.021 | NA | 0.000 | 1.257 | 1.166 |
| CA10 |  | 6 | 0.464 | 0.002 | 6 | 0.59 | 8 | 0.931 | 6 | 0.014 | NA | 0.000 | 1.523 | -1.059 |
| **Cayo Arenas** | | 84 | 3.536 | 0.107 | 84 | 0.079 | 112 | 1.547 | 84 | 0.37 | 84 | 1.0 | 2.774 | 0.762 |

Shaded cells are sites that did not complete the minimum number of transects during the surveys. For transects without information, the site mean was assigned. *For Cayo Arenas 15, the nighttime survey was conducted in 2024, one year after daytime monitoring.

**Table S3**. Modified coral density averages and linear extension rates for Cayo Arenas, the Campeche Bank, and the Gulf of Mexico.

| **Species** | **linear growth (cm/yr)** | **skeletal density (g/cm^3^)** |
| --- | --- | --- |
| *Acropora cervicornis* | 4** | 1.96*** |
| *Diploria labyrinthiformis* | 0.35** | 1.43*** |
| *Millepora complanata* | 0.8** | 1.51*** |
| *Montastraea cavernosa* | 0.32** | 1.64*** |
| *Orbicella annularis* | 0.86° | 1.73° |
| *Orbicella faveolata* | 0.91** | 1.42** |
| *Orbicella faveolata* | 0.82° | 1.04° |
| *Porites astreoides* | 0.51** | 1.48** |
| *Porites porites* | 1.92** | 1.2*** |
| *Porites furcata* | 1.76** | 1.05*** |
| *Pseudodiploria clivosa* | 0.48** | 1.2*** |
| *Pseudodiploria strigosa* | 0.57** | 1.21** |
| *Siderastrea radians* | 0.19** | 1.51*** |
| *Siderastrea siderea* | 0.38** | 1.41** |

Skeletal density values and linear growth rates for Cayo Arenas° (Carricart-Ganivet and Merino, 2001); Gulf of Mexico **(Vaughn, 1915; Hudson et al 1994; Witman 1988; Weber and White 1977; Manzello et al. 2015; Kissling, 1977; Elizalde-Rendon et al, 2010; Castillo, 2018; Manzello et al 2021; Rezak et al 1985; Horta-Puga et al. 2014) and the Caribbean Sea*** (means of ReefBudget metodology database, Perry and Lange, 2019).

**Table S4**. Classification of Mexican Caribbean sites.

| **Managed Fishery** | **Zone** | **Site** | **Year** | **Reference** |
| --- | --- | --- | --- | --- |
| Well-Managed Fisheries | Cancun | Cuevones | 2019 | Espinosa-Andrade et al., 2020; Perry et al., 2025; Molina-Hernández et al., 2020 |
| Well-Managed Fisheries | Cancun | Manchones | 2019 | Perry et al., 2025 |
| Well-Managed Fisheries | Cancun | Nizuc 5m | 2019 | Perry et al., 2025 |
| Well-Managed Fisheries | Cancun | Nizuc C3 | 2018 | Perry et al., 2025 |
| Well-Managed Fisheries | Cancun | Nizuc F3 | 2018 | Perry et al., 2025 |
| Well-Managed Fisheries | Puerto Morelos | Bocana Caricomp | 2018 | Perry et al., 2025 |
| Well-Managed Fisheries | Puerto Morelos | Bonanza | 2017 | Perry et al., 2025 |
| Well-Managed Fisheries | Puerto Morelos | Bonanza Profundo | 2018, 2019 | Perry et al., 2025 |
| Well-Managed Fisheries | Puerto Morelos | La catedral 5m | 2019 | Perry et al., 2025 |
| Well-Managed Fisheries | Puerto Morelos | La catedral Posterior | 2019 | Espinosa-Andrade et al., 2020; Perry et al., 2025 |
| Well-Managed Fisheries | Puerto Morelos | Limones | 2017 | Espinosa-Andrade et al., 2020; Perry et al., 2025 |
| Well-Managed Fisheries | Puerto Morelos | Manchones Norte | 2018, 2019 | Espinosa-Andrade et al., 2020; Perry et al., 2025 |
| Well-Managed Fisheries | Puerto Morelos | Mar F2 | 2018 | Espinosa-Andrade et al., 2020; Perry et al., 2025 |
| Well-Managed Fisheries | Puerto Morelos | Mar F4 | 2018 | Espinosa-Andrade et al., 2020; Perry et al., 2025 |
| Well-Managed Fisheries | Puerto Morelos | PM-F2 | 2018 | Perry et al., 2025 |
| Well-Managed Fisheries | Puerto Morelos | Radio Pirata | 2018 | Espinosa-Andrade et al., 2020; Perry et al., 2025 |
| Well-Managed Fisheries | Puerto Morelos | Tanchacte | 2018 | Espinosa-Andrade et al., 2020; Perry et al., 2025;  Molina-Hernández et al. 2020 |
| Well-Managed Fisheries | Cozumel | Caracolillo | 2018 | Perry et al., 2025 |
| Well-Managed Fisheries | Cozumel | Chankanaab | 2017 | Espinosa-Andrade et al., 2020; Perry et al., 2025;  Molina-Hernández et al. 2020 |
| Well-Managed Fisheries | Cozumel | Colombia Somero | 2018 | Perry et al., 2025 |
| Well-Managed Fisheries | Cozumel | Hanan | 2018, 2019 | Espinosa-Andrade et al., 2020; Perry et al., 2025 |
| Well-Managed Fisheries | Cozumel | Microatolones 5m | 2019 | Perry et al., 2025 |
| Well-Managed Fisheries | Cozumel | MX3054 | 2018, 2019 | Molina-Hernández et al., 2020; Perry et al., 2025 |
| Well-Managed Fisheries | Cozumel | Palancar Jardines | 2017 | Perry et al., 2025 |
| Well-Managed Fisheries | Cozumel | Paraiso | 2017 | Espinosa-Andrade et al., 2020; Perry et al., 2025 |
| Well-Managed Fisheries | Cozumel | Punta Sur Somero B | 2018 | Espinosa-Andrade et al., 2020; Perry et al., 2025; Molina-Hernández et al., 2020; |
| Under-Managed Fisheries | Punta Maroma | Mar F5 | 2019, 2021 | Perry et al., 2025 |
| Under-Managed Fisheries | Punta Maroma | Mar P1 | 2019 | Perry et al., 2025 |
| Under-Managed Fisheries | Punta Maroma | Punta Maroma Norte | 2019, 2021 | Perry et al., 2025 |
| Under-Managed Fisheries | Punta Maroma | Punta Maroma Sur | 2019, 2021 | Perry et al., 2025 |
| Under-Managed Fisheries | Akumal | Akumal Langosta | 2021 | Perry et al., 2025 |
| Under-Managed Fisheries | Akumal | Dicks | 2018, 2021 | Perry et al., 2025 |
| Under-Managed Fisheries | Akumal | Media Luna | 2018, 2019 | Perry et al., 2025 |
| Under-Managed Fisheries | Akumal | Yalku | 2018, 2021 | Perry et al., 2025 |
| Under-Managed Fisheries | Tulum | Casa Cenote | 2021 | Espinosa-Andrade et al., 2020 |
| Under-Managed Fisheries | Tulum | Casa Cenote Profundo | 2021 | Espinosa-Andrade et al., 2020 |
| Under-Managed Fisheries | Punta Allen | MX1006 | 2021 | Perry et al., 2025 |
| Under-Managed Fisheries | Punta Allen | MX2007 | 2021 | Perry et al., 2025 |
| Under-Managed Fisheries | Punta Allen | Niccehabin 6m | 2018, 2021 | Perry et al., 2025 |
| Under-Managed Fisheries | Punta Allen | Niccehabin Frontal Somero | 2018 | Perry et al., 2025 |
| Under-Managed Fisheries | Punta Allen | Niccehabin Posterior | 2018 | Perry et al., 2025 |
| Under-Managed Fisheries | Punta Allen | Punta Allen Centro | 2018, 2021 | Perry et al., 2025 |
| Under-Managed Fisheries | Punta Allen | Punta Allen Norte | 2018, 2021 | Perry et al., 2025 |
| Under-Managed Fisheries | Punta Allen | San Antonio 6m | 2018, 2021 | Perry et al., 2025 |
| Under-Managed Fisheries | Punta Allen | San Antonio Posterior | 2018 | Perry et al., 2025 |
| Under-Managed Fisheries | Punta Allen | Yuyum 6m | 2018 | Perry et al., 2025 |
| Under-Managed Fisheries | Punta Allen | Yuyum Frontal Somero | 2018 | Perry et al., 2025 |
| Under-Managed Fisheries | Punta Allen | Yuyum Posterior | 2018 | Perry et al., 2025 |
| Under-Managed Fisheries | Mahahual | 40 Cañones | 2017 | Perry et al., 2018 |
| Under-Managed Fisheries | Mahahual | El Faro | 2017 | Perry et al., 2018; 2025 |
| Under-Managed Fisheries | Mahahual | Faro Viejo | 2017 | Perry et al., 2018, 2025 |
| Under-Managed Fisheries | Mahahual | Hotel Arenas | 2017 | Perry et al., 2018, 2025 |
| Under-Managed Fisheries | Mahahual | Mahahual | 2017, 2019 | Perry et al., 2018 |
| Under-Managed Fisheries | Mahahual | Mahahual Centro | 2017, 2019 | Perry et al., 2018, 2025 |
| Under-Managed Fisheries | Mahahual | MX1020 | 2017 | Perry et al., 2008; Molina-Hernández et al., 2020 |
| Under-Managed Fisheries | Mahahual | Puerto Angel | 2017 | Perry et al., 2018; Molina-Hernández et al., 2020 |
| Under-Managed Fisheries | Mahahual | Quebrado 5m | 2019 | Perry et al., 2018 |
| Under-Managed Fisheries | Xcalak | Bacalar Chico | 2019 | Espinosa-Andrade et al., 2020 |
| Under-Managed Fisheries | Xcalak | Doña Nica | 2019 | Espinosa-Andrade et al., 2020 |
| Under-Managed Fisheries | Xcalak | Portillas | 2019 | Perry et al., 2025 |
| Under-Managed Fisheries | Xcalak | Portillas 5m | 2019 | Perry et al., 2025 |
| Under-Managed Fisheries | Xcalak | Portillas Cresta | 2019 | Perry et al., 2025 |
| Under-Managed Fisheries | Xcalak | Xahuayxol | 2019 | Espinosa-Andrade et al., 2020 |
| Under-Managed Fisheries | Isla Contoy | Ixlaché | 2017 | Perry et al., 2025 |
| Under-Managed Fisheries | Banco Chinchorro | CHI02 | 2019 | Espinosa-Andrade et al., 2020 |
| Under-Managed Fisheries | Banco Chinchorro | CHI04 | 2019 | Perry et al., 2025 |
| Under-Managed Fisheries | Banco Chinchorro | CHI05 | 2019 | Perry et al., 2025 |
| Under-Managed Fisheries | Banco Chinchorro | Far Star | 2019 | Perry et al., 2025 |
| Under-Managed Fisheries | Banco Chinchorro | Isla che | 2019 | Perry et al., 2025 |
| Under-Managed Fisheries | Banco Chinchorro | La Baliza 2 | 2019 | Perry et al., 2025 |
| Under-Managed Fisheries | Banco Chinchorro | MX3082 | 2019 | Perry et al., 2025 |

**Table S5**. Parrotfish bioerosion rates (kg CaCO_3_ m^2^ yr^-1^±SE) in Cayo Arenas.

| **Zone** | **Site** | **Transects** | ***Sparisoma viride*** | | ***Scarus vetula*** | | ***Scarus taeniopterus*** | | ***Scarus***  ***iseri*** | | ***Scarus***  ***coeruleus*** | | ***Sparisoma aurofrenatum*** | | ***Sparisoma rubripinne*** | | ***Sparisoma chrysopterum*** | |
| --- | --- | --- | --- | --- | --- | --- | --- | --- | --- | --- | --- | --- | --- | --- | --- | --- | --- | --- |
|  |  |  | **Mean** | **SE** | **Mean** | **SE** | **Mean** | **SE** | **Mean** | **SE** | **Mean** | **SE** | **Mean** | **SE** | **Mean** | **SE** | **Mean** | **SE** |
| Exposed | CA15 | 8 | 2.53 | 0.88 | 1.29 | 0.27 | 0.07 | 0.04 | 0.00 | 0.00 | 0 | 0 | 0.00 | 0.00 | 0.00 | 0.00 | 0 | 0 |
| Exposed | CA11 | 8 | 1.03 | 0.69 | 0.09 | 0.09 | 0.10 | 0.04 | 0.00 | 0.00 | 0 | 0 | 0.01 | 0.00 | 0.00 | 0.00 | 0 | 0 |
| Exposed | CA5 | 9 | 1.92 | 0.68 | 0.17 | 0.15 | 0.13 | 0.05 | 0.00 | 0.00 | 0 | 0 | 0.02 | 0.00 | 0.01 | 0.01 | 0 | 0 |
| Exposed | CA16 | 8 | 2.27 | 1.10 | 0.05 | 0.03 | 0.07 | 0.03 | 0.12 | 0.05 | 0 | 0 | 0.01 | 0.00 | 0.01 | 0.01 | 0 | 0 |
| Exposed | Cayo Arenas Acropora | 8 | 1.27 | 0.83 | 0.51 | 0.15 | 0.06 | 0.04 | 0.03 | 0.01 | 0 | 0 | 0.01 | 0.00 | 0.00 | 0.00 | 0 | 0 |
| Protected | CA6 | 10 | 1.58 | 0.80 | 0.02 | 0.02 | 0.04 | 0.03 | 0.01 | 0.00 | 0.0028 | 0.0028 | 0.02 | 0.00 | 0.00 | 0.00 | 0 | 0 |
| Protected | CA1 | 8 | 1.17 | 0.38 | 0.10 | 0.04 | 0.03 | 0.01 | 0.03 | 0.01 | 0 | 0 | 0.02 | 0.01 | 0.00 | 0.00 | 0 | 0 |
| Protected | CA13 | 7 | 1.83 | 0.54 | 0.03 | 0.03 | 0.06 | 0.06 | 0.02 | 0.01 | 0 | 0 | 0.01 | 0.00 | 0.00 | 0.00 | 0 | 0 |
| Bajos | Bajo Lucas | 8 | 0.49 | 0.27 | 0.00 | 0.00 | 0.03 | 0.02 | 0.01 | 0.01 | 0 | 0 | 0.01 | 0.00 | 0.00 | 0.00 | 0.0007 | 0.0004 |
| Bajos | Bajo tortugas | 8 | 0.35 | 0.24 | 0.09 | 0.08 | 0.02 | 0.02 | 0.02 | 0.01 | 0 | 0 | 0.01 | 0.00 | 0.00 | 0.00 | 0 | 0 |
| Lagoon | CA14 | 7 | 0.37 | 0.25 | 0.00 | 0.00 | 0.00 | 0.00 | 0.05 | 0.02 | 0.002499284 | 0.002499284 | 0.01 | 0.00 | 0.00 | 0.00 | 0 | 0 |
| Lagoon | CA8 | 7 | 1.07 | 0.38 | 0.00 | 0.00 | 0.05 | 0.02 | 0.00 | 0.00 | 0 | 0 | 0.02 | 0.01 | 0.00 | 0.00 | 0 | 0 |
| Lagoon | CA4 | 7 | 0.94 | 0.47 | 0.19 | 0.19 | 0.00 | 0.00 | 0.04 | 0.02 | 0 | 0 | 0.02 | 0.00 | 0.00 | 0.00 | 0 | 0 |
| Lagoon | CA10 | 8 | 0.88 | 0.42 | 0.00 | 0.00 | 0.00 | 0.00 | 0.03 | 0.01 | 0 | 0 | 0.01 | 0.00 | 0.00 | 0.00 | 0.0004 | 0.00043 |
| Cayo Arenas | | 111 | 1.28 | 0.17 | 0.18 | 0.04 | 0.05 | 0.01 | 0.03 | 0.00 | 0.0004 | 0.0003 | 0.01 | 0.00 | 0.00 | 0.00 | 8.19354E-05 | 4.61276E-05 |

**Table S6.** Sea urchin nighttime bioerosion rates (kg CaCO_3_ m^2^ yr^-1^±SE) in Cayo Arenas.

| **Zone** | **Site** | **Transects** | ***Diadema antillarum*** | | ***Echinometra viridis*** | | ***Echinometra tribuloides*** | | ***Eucidaris lucunter*** | |
| --- | --- | --- | --- | --- | --- | --- | --- | --- | --- | --- |
|  |  |  | **Mean** | **SE** | **Mean** | **SE** | **Mean** | **SE** | **Mean** | **SE** |
| Exposed | CA15 | 7 | 1.58 | 1.48 | 0.02 | 0.01 | 0.00 | 0.01 | 0.00 | 0.00 |
| Protected | CA 1 | 7 | 1.09 | 0.13 | 0.09 | 0.03 | 0.00 | 0.00 | 0.01 | 0.01 |
| Bajos | Bajo Tortugas | 5 | 3.85 | 0.02 | 0.05 | 0.00 | 0.01 | 0.00 | 0.00 | 0.00 |
| Lagoon | CA14 | 6 | 0.02 | 0.22 | 0.00 | 0.01 | 0.00 | 0.00 | 0.00 | 0.00 |
| Cayo Arenas | | 25 | 1.52 | 0.38 | 0.04 | 0.01 | 0.00 | 0.00 | 0.00 | 0.00 |

**Table S7**. Coral cover (%±SE) of Cayo Arenas. Classification based on morphology coral.

| **Zone** | **Site** | **Transects** | ***Orbicella* spp.** | | **Massive others** | | **Submassive** | | **Branching** | | **Digitate** | | ***Millepora* spp.** | | **No framework corals** | | **Total coral cover** | | **CCA** | |
| --- | --- | --- | --- | --- | --- | --- | --- | --- | --- | --- | --- | --- | --- | --- | --- | --- | --- | --- | --- | --- |
|  |  |  | **Mean** | **SE** | **Mean** | **SE** | **Mean** | **SE** | **Mean** | **SE** | **Mean** | **SE** | **Mean** | **SE** | **Mean** | **SE** | **Mean** | **SE** | **Mean** | **SE** |
| **Exposed** | CA11 | 6 | 5.67 | 3.15 | 1.94 | 4.40 | 1.64 | 5.38 | 0.00 |  | 8.11 | 1.54 | 8.88 | 2.53 | 0.28 | 8.19 | 26.52 | 1.58 | 31.56 | 1.24 |
|  | CA15 | 6 | 9.01 | 2.13 | 1.42 | 3.36 | 2.51 | 4.78 | 0.00 |  | 0.33 | 6.69 | 9.13 | 2.38 | 0.25 | 9.60 | 22.65 | 1.90 | 31.91 | 1.44 |
|  | CA5 | 6 | 2.52 | 2.93 | 3.89 | 2.85 | 0.08 | 13.68 | 3.00 | 3.08 | 0.00 |  | 4.53 | 2.35 | 6.80 | 1.94 | 20.81 | 1.99 | 10.02 | 2.33 |
|  | CA16 | 6 | 5.01 | 2.38 | 9.26 | 4.90 | 0.89 | 7.83 | 0.00 |  | 0.00 |  | 0.59 | 7.21 | 0.13 | 10.72 | 15.88 | 2.42 | 10.08 | 2.17 |
|  | Cayo  Arenas Acropora | 6 | 1.29 | 3.38 | 1.39 | 4.61 | 4.01 | 5.89 | 7.84 | 2.15 | 0.00 |  | 18.04 | 1.76 | 0.05 | 16.57 | 32.63 | 1.51 | 18.41 | 2.02 |
| **Protected** | CA6 | 6 | 20.39 | 2.49 | 4.98 | 3.47 | 0.04 | 18.29 | 0.00 |  | 0.00 |  | 0.63 | 5.88 | 0.72 | 6.51 | 26.77 | 2.69 | 4.69 | 2.55 |
|  | CA1 | 6 | 21.73 | 2.16 | 11.20 | 2.11 | 1.69 | 5.53 | 0.00 |  | 0.00 |  | 0.17 | 9.28 | 0.79 | 6.70 | 35.58 | 1.91 | 3.73 | 3.27 |
|  | CA13 | 6 | 5.43 | 2.10 | 0.00 |  | 0.00 |  | 0.00 |  | 39.64 | 1.45 | 0.22 | 8.18 | 0.16 | 11.51 | 45.45 | 1.36 | 3.72 | 2.99 |
| **Bajos** | Bajo Tortugas | 6 | 15.62 | 1.77 | 10.21 | 2.55 | 1.48 | 3.71 | 0.00 |  | 0.75 | 6.40 | 1.31 | 4.10 | 0.99 | 4.25 | 30.37 | 2.11 | 16.65 | 1.91 |
|  | Bajo  Lucas | 6 | 27.47 | 2.02 | 6.17 | 2.74 | 1.21 | 5.14 | 0.00 |  | 1.52 | 5.43 | 0.51 | 7.17 | 2.34 | 4.26 | 39.22 | 2.28 | 14.89 | 2.35 |
| **Lagoon** | CA14 | 6 | 14.15 | 1.94 | 12.79 | 1.82 | 1.43 | 4.70 | 0.00 |  | 0.00 |  | 0.41 | 7.13 | 0.29 | 8.13 | 29.06 | 2.08 | 6.43 | 3.46 |
|  | CA8 | 6 | 22.43 | 1.60 | 4.81 | 3.17 | 0.33 | 7.31 | 0.00 |  | 0.04 | 19.59 | 0.00 |  | 0.19 | 8.81 | 27.79 | 1.64 | 8.32 | 2.90 |
|  | CA4 | 5 | 12.07 | 1.87 | 5.70 | 2.44 | 1.06 | 5.46 | 0.00 |  | 0.28 | 9.52 | 0.00 |  | 0.18 | 11.82 | 19.29 | 2.03 | 10.90 | 2.40 |
|  | CA10 | 6 | 29.17 | 2.37 | 8.67 | 2.51 | 1.00 | 4.24 | 0.00 |  | 0.00 |  | 0.33 | 8.35 | 0.17 | 9.39 | 39.33 | 2.18 | 2.83 | 3.43 |
| Cayo Arenas | | 83 | 13.73 | 1.75 | 5.89 | 2.43 | 1.24 | 4.74 | 0.78 | 3.45 | 3.66 | 1.75 | 3.24 | 2.35 | 0.96 | 3.44 | 29.50 | 1.67 | 12.46 | 1.67 |

**Table S8**. Population density (ind m^-2^ ±SE) of sea urchins at Cayo Arenas.

| **Reef System** | **Managed Fishery** | **Zone** | **Site** | **Years** | **Transects** | ***D. antillarum*** | | ***Echinometra* spp.** | | ***Eucidaris tribuloides*** | |
| --- | --- | --- | --- | --- | --- | --- | --- | --- | --- | --- | --- |
|  |  |  |  |  |  | **Mean** | **SE** | **Mean** | **SE** | **Mean** | **SE** |
| **Nighttime surveys** | | | | | | | | | | | |
| Cayo Arenas | Remote reef | Exposed | CA15 | 2024 | 22 | 1.05 | 0.17 | 0.1 | 0.03 | 0.018181818 | 0.01 |
| Cayo Arenas | Remote reef | Protected | CA1 | 2023 | 7 | 0.842857143 | 0.10 | 0.8 | 0.22 | 0.042857143 | 0.04 |
| Cayo Arenas | Remote reef | Bajos | Bajo Tortugas | 2023 | 5 | 2.46 | 0.90 | 0.36 | 0.09 | 0.02 | 0.02 |
| Cayo Arenas | Remote reef | Lagoon | CA14 | 2023 | 6 | 0.016666667 | 0.02 | NA | 0.00 | NA | 0.00 |
| **Daytime surveys** | | | | | | | | | | | |
| Cayo Arenas | Remote reef | Exposed | Cayo Arenas Acropora |  | 6 | 0.13 | 0.02 | 0.12 | 0.05 | 0.12 | 0.05 |
| Cayo Arenas | Remote reef | Exposed | CA11 |  | 6 | 0.87 | 0.14 | 0.03 | 0.02 | NA | 0.00 |
| Cayo Arenas | Remote reef | Exposed | CA15 |  | 6 | 0.72 | 0.22 | 0.08 | 0.08 | NA | 0.00 |
| Cayo Arenas | Remote reef | Exposed | Ca16 |  | 6 | 0.17 | 0.08 | 0.08 | 0.05 | 0.02 | 0.02 |
| Cayo Arenas | Remote reef | Exposed | CA5 |  | 6 | 0.37 | 0.09 | NA | 0.00 | NA | 0.00 |
| Cayo Arenas | Remote reef | Protected | CA1 |  | 5 | 0.22 | 0.10 | 0.34 | 0.09 | NA | 0.00 |
| Cayo Arenas | Remote reef | Protected | CA13 |  | 6 | 0.05 | 0.03 | NA | 0.00 | NA | 0.00 |
| Cayo Arenas | Remote reef | Protected | CA6 |  | 6 | 0.13 | 0.06 | 0.07 | 0.04 | NA | 0.00 |
| Cayo Arenas | Remote reef | Bajos | Bajo Lucas |  | 6 | 0.57 | 0.11 | 0.28 | 0.07 | NA | 0.00 |
| Cayo Arenas | Remote reef | Bajos | Bajo Tortugas |  | 6 | 0.22 | 0.07 | 0.05 | 0.03 | NA | 0.00 |
| Cayo Arenas | Remote reef | Lagoon | CA10 |  | 5 | NA | 0.00 | 0.04 | 0.04 | 0.10 | 0.04 |
| Cayo Arenas | Remote reef | Lagoon | CA14 |  | 6 | NA | 0.00 | NA | 0.00 | NA | 0.00 |
| Cayo Arenas | Remote reef | Lagoon | CA4 |  | 6 | 0.02 | 0.02 | 0.07 | 0.03 | NA | 0.00 |
| Cayo Arenas | Remote reef | Lagoon | CA8 |  | 6 | 0.02 | 0.02 | NA | 0.00 | NA | 0.00 |

**Table S9**. Endolithic sponges bioerosion rates (kg CaCO_3_ m^2^ yr^-1^±SE) in Cayo Arenas.

| **Zone** | **Site** | **Transects** | ***Cliona caribbaea*** | | ***Cliona***  ***delitrix*** | | ***Cliona***  ***tenuis*** | | ***Cliona***  ***aprica*** | | ***Siphonodictyon***  ***spp*** | | ***Cliona***  ***varians*** | | **Total** | |
| --- | --- | --- | --- | --- | --- | --- | --- | --- | --- | --- | --- | --- | --- | --- | --- | --- |
|  |  |  | **Mean** | **SE** | **Mean** | **SE** | **Mean** | **SE** | **Mean** | **SE** | **Mean** | **SE** | **Mean** | **SE** | **Mean** | **SE** |
| Exposed | CA11 | 6 | 0.01 | 0.01 | 0.00 | 0.00 | 0.03 | 0.03 | 0.00 | 0.00 | 0.00 | 0.00 | 0.00 | 0.00 | 0.04 | 0.03 |
| Exposed | CA15 | 6 | 0.02 | 0.01 | 0.05 | 0.02 | 0.00 | 0.00 | 0.00 | 0.00 | 0.00 | 0.00 | 0.00 | 0.00 | 0.07 | 0.02 |
| Exposed | CA16 | 6 | 0.00 | 0.00 | 0.02 | 0.01 | 0.00 | 0.00 | 0.00 | 0.00 | 0.00 | 0.00 | 0.00 | 0.00 | 0.02 | 0.01 |
| Exposed | CA5 | 6 | 0.00 | 0.00 | 0.00 | 0.00 | 0.00 | 0.00 | 0.00 | 0.00 | 0.00 | 0.00 | 0.00 | 0.00 | 0.00 | 0.00 |
| Exposed | Cayo Arenas Acropora | 6 | 0.01 | 0.01 | 0.02 | 0.02 | 0.13 | 0.06 | 0.00 | 0.00 | 0.01 | 0.00 | 0.00 | 0.00 | 0.17 | 0.06 |
| Protected | CA6 | 6 | 0.00 | 0.00 | 0.00 | 0.00 | 0.03 | 0.02 | 0.00 | 0.00 | 0.00 | 0.00 | 0.00 | 0.00 | 0.03 | 0.02 |
| Protected | CA1 | 5 | 0.01 | 0.00 | 0.01 | 0.00 | 0.00 | 0.00 | 0.00 | 0.00 | 0.00 | 0.00 | 0.00 | 0.00 | 0.02 | 0.00 |
| Protected | CA13 | 6 | 0.00 | 0.00 | 0.00 | 0.00 | 0.00 | 0.00 | 0.00 | 0.00 | 0.00 | 0.00 | 0.00 | 0.00 | 0.00 | 0.00 |
| Bajos | Bajo Tortugas | 6 | 0.01 | 0.00 | 0.05 | 0.01 | 0.00 | 0.00 | 0.00 | 0.00 | 0.00 | 0.00 | 0.00 | 0.00 | 0.06 | 0.01 |
| Bajos | Bajo Lucas | 6 | 0.02 | 0.01 | 0.01 | 0.01 | 0.01 | 0.01 | 0.00 | 0.00 | 0.01 | 0.01 | 0.01 | 0.01 | 0.05 | 0.01 |
| Lagoon | CA14 | 6 | 0.01 | 0.01 | 0.02 | 0.01 | 0.00 | 0.00 | 0.00 | 0.00 | 0.00 | 0.00 | 0.00 | 0.00 | 0.02 | 0.01 |
| Laguna | CA8 | 6 | 0.00 | 0.00 | 0.00 | 0.00 | 0.00 | 0.00 | 0.00 | 0.00 | 0.00 | 0.00 | 0.00 | 0.00 | 0.00 | 0.00 |
| Laguna | CA4 | 6 | 0.01 | 0.00 | 0.02 | 0.01 | 0.00 | 0.00 | 0.00 | 0.00 | 0.00 | 0.00 | 0.00 | 0.00 | 0.03 | 0.01 |
| Laguna | CA10 | 5 | 0.32 | 0.09 | 0.24 | 0.05 | 0.00 | 0.00 | 0.03 | 0.03 | 0.00 | 0.00 | 0.00 | 0.00 | 0.59 | 0.08 |
| Cayo Arenas | | 82 | 0.03 | 0.01 | 0.03 | 0.01 | 0.01 | 0.01 | 0.00 | 0.00 | 0.00 | 0.00 | 0.00 | 0.00 | 0.07 | 0.02 |

**Table S10**. Microbioerosion rates (kg CaCO_3_ m^2^ yr^-1^±SE) in Cayo Arenas.

| **Zone** | **Sites** | **Transects** | **Microbioerosion** | |
| --- | --- | --- | --- | --- |
|  |  |  | **Mean** | **SE** |
| Exposed | CA11 | 6 | 0.11832 | 0.029457756 |
| Exposed | CA15 | 6 | 0.11816 | 0.027486142 |
| Exposed | CA16 | 6 | 0.11584 | 0.021100688 |
| Exposed | CA5 | 6 | 0.15512 | 0.021972667 |
| Exposed | Cayo Arenas Acropora | 6 | 0.06944 | 0.015523958 |
| Protected | CA6 | 6 | 0.15984 | 0.016422502 |
| Protected | CA1 | 6 | 0.1816 | 0.032733259 |
| Protected | CA13 | 6 | 0.1272 | 0.021320863 |
| Bajos | Bajo Tortugas | 6 | 0.091702222 | 0.019350335 |
| Bajos | Bajo Lucas | 6 | 0.1126 | 0.018341027 |
| Lagoon | CA14 | 6 | 0.10748 | 0.026442178 |
| Lagoon | CA8 | 6 | 0.09088 | 0.011052804 |
| Lagoon | CA4 | 5 | 0.034752 | 0.015605058 |
| Lagoon | CA10 | 6 | 0.00196 | 0.000190158 |
| Cayo Arenas | | 83 | 0.11 | 0.01 |
